# Scikit-NeuroMSI: A Generalized Framework for Modeling Multisensory Integration

**DOI:** 10.1101/2025.05.26.656124

**Authors:** Renato Paredes, Juan B. Cabral, Peggy Seriès

**Affiliations:** Departament of Psychology, Pontifical Catholic University of Peru, Lima, Peru; Instituto de Investigaciones Psicológicas, Facultad de Psicología, Universidad Nacional de Córdoba, Córdoba, Argentina; Grupo de Innovación y Desarrollo Tecnológico, Gerencia De Vinculación Tecnológica, Centro Espacial Teófilo Tabanera, Comisión Nacional de Actividades Espaciales (CONAE), Córdoba, Argentina; Consejo Nacional de Investigaciones Científicas y Técnicas (CONICET), Buenos Aires, Argentina; School of Informatics, University of Edinburgh, Edinburgh, United Kingdom

**Keywords:** Multisensory Integration, Causal Inference, Scientific Software, Computational Neuroscience, Computational Models

## Abstract

Multisensory integration is a fundamental neural mechanism crucial for understanding cognition. Multiple theoretical models exist to account for the computational processes underpinning this mechanism. However, there is an absence of a consolidated framework that facilitates the examination of multisensory integration across diverse experimental and computational contexts. We introduce Scikit-NeuroMSI, an accessible Python-based open-source framework designed to streamline the implementation and evaluation of computational models of multisensory integration. The capabilities of Scikit-NeuroMSI were demonstrated in enabling the implementation of multiple models of multisensory integration at different levels of analysis. Furthermore, we illustrate the utility of the software in systematically exploring the model’s behavior in spatiotemporal causal inference tasks through parameter sweeps in simulations. Particularly, we conducted a comparative analysis of Bayesian and network models of multisensory integration to identify commonalities that may enable to bridge both levels of description, addressing a key research question within the field. We discuss the significance of this approach in generating computationally informed hypotheses in multisensory research. Recommendations for the improvement of this software and directions for future research using this framework are presented.

## 1 Introduction

Multisensory integration is the neural process by which signals originating from distinct sensory modalities (such as visual, tactile, or auditory) are merged. As a result, the multisensory response can differ significantly from the responses elicited by stimuli confined to a single sensory modality (Stein et al., 2010; Stein & Stanford, 2008). Disturbances in multisensory processing can impact various cognitive domains (Wallace, Woynaroski, & Stevenson, 2020). For example, alterations in multisensory function are observed in various neuropsychiatric and neurological disorders (e.g. SCZ, ASD, dementia, sensory loss, dyslexia) (Cascio, Foss-Feig, Burnette, Heacock, & Cosby, 2012; Festa, Katz, Ott, Tremont, & Heindel, 2017; Hahn, Foxe, & Molholm, 2014; Haß et al., 2017; B. Martin, Giersch, Huron, & van Wassenhove, 2013; Noel, Paredes, et al., 2022; Paredes, Ferri, & Seriès, 2022; Ramkhalawansingh, Keshavarz, Haycock, Shahab, & Campos, 2017; Stevenson et al., 2014; Wu et al., 2012; Zhou et al., 2018; Zvyagintsev, Parisi, & Mathiak, 2017). Furthermore, a substantial body of research is exploring the potential of multisensory markers to predict future clinical manifestations or to serve as key focal points for therapeutic interventions (Bolognini, Rasi, Coccia, & Làdavas, 2005; Gieseler, Tahden, Thiel, & Colonius, 2018; Sánchez, Millán-Calenti, Lorenzo-López, & Maseda, 2013).

Computational modeling is crucial for the advancement of the field due to its potential to develop formal theories of the neural mechanisms of multisensory integration (Colonius & Diederich, 2020; Meijer & Noppeney, 2020). It forces scientists to analyze, specify, and formalize their ideas, while also allowing them to generate precise quantitative predictions suitable for testing in future experiments (Blohm, Kording, & Schrater, 2020; Guest & Martin, 2021). Consequently, the body of literature on computational models of multisensory integration has experienced substantial growth over the past two decades (Colonius & Diederich, 2020).

The main modeling approaches are optimal cue combination (Alais & Burr, 2004; Ernst & Banks, 2002; Fetsch, DeAngelis, & Angelaki, 2013; Parise & Ernst, 2016), Bayesian causal inference (Körding et al., 2007; Meijer & Noppeney, 2020; Rohe, Ehlis, & Noppeney, 2019; Rohe & Noppeney, 2015; Shams & Beierholm, 2010), race (Colonius & Diederich, 2004; Colonius, Wolff, & Diederich, 2017; Diederich, 1992, 1995), and network (Cuppini, Shams, Magosso, & Ursino, 2017; Ma & Pouget, 2008; Ma & Rahmati, 2013; Miller, Stein, & Rowland, 2017; Ohshiro, Angelaki, & DeAngelis, 2011; Ursino, Cuppini, Magosso, Beierholm, & Shams, 2019) models. Despite being based on very general mechanisms, these models are typically restricted to a specific experimental paradigm (e.g. spatial localization, orientation judgments, temporal order judgments, rate detection, sound-induced flash illusion, among others) and vary in their level of description (e.g. computation, implementation or algorithm) and sophistication.

Overall, the field lacks a unified theoretical approach to multisensory integration that allows for model testing in different experimental and computational paradigms. There is a growing need for scientific software specifically designed to represent the distinctive concepts and mechanisms that characterize the integration of information from different sensory modalities. To our knowledge, there is as yet no software that can provide a unified computational environment designed to facilitate the examination of differences in model predictions. Multisensory integration modelers currently rely on packages built for general-purpose computational neuroscience modeling, such as Brian (Stimberg, Brette, & Goodman, 2019), HDDM (Wiecki, Sofer, & Frank, 2013), TAPAS (Frässle et al., 2021) or PyRates (Gast et al., 2019), BrainPy (Wang et al., 2023), among others. There are also alternatives for specific multisensory integration models, such as the Bayesian Causal Inference Toolbox (Zhu, Beierholm, & Shams, 2024a), but no frameworks encompassing more than one modeling approach.

Here we present Scikit-NeuroMSI (Paredes, Seriès, & Cabral, 2023), an open-source Python (Rossum & Drake, 2010) framework that simplifies the implementation of neurocomputational models of multisensory integration. The package currently allows to run seminal computational models of multisensory integration and to easily implement new models defined by users. As an illustration, we show how this framework facilitates the analysis of spatiotemporal multisensory integration at different levels of description using Bayesian and network models.

This paper targets a diverse audience in neuroscience and computational fields, including computational neuroscientists and researchers in sensory processing. It is also beneficial for software developers and engineers interested in multisensory integration models. While basic neuroscience and programming knowledge is useful, proficiency in Python is needed to understand the implementation of Scikit-NeuroMSI, as it is developed in this language. We make the paper accessible by covering theoretical foundations and practical implementations, with explanations and code examples to aid comprehension. Those with Python skills can dive directly into the implementation details, while newcomers will find enough context to grasp the concepts. This approach highlights the challenge of standardizing multisensory integration models, which requires a blend of neuroscience, mathematics, and software engineering.

## 2 Software

### 2.1 Design overview

Experiments on multisensory integration investigate how the brain combines information from multiple sensory modalities to create a unified perception of the environment. These studies often involve manipulating the reliability or congruency of sensory cues from different modalities, such as vision and touch, to examine how the brain weighs and integrates this information (Colonius & Diederich, 2020). Experimental paradigms include spatial localisation tasks for cross-modal spatial interactions (Alais & Burr, 2004; Ernst & Banks, 2002), temporal order judgments for timing perception of multisensory events (Ferri, Ambrosini, & Costantini, 2016), or the double flash illusion for auditory influence on visual perception (Shams, Kamitani, & Shimojo, 2002), among others.

Computational models play a crucial role in multisensory research by providing frameworks to predict and explain behavioral and neural responses in multisensory contexts (Chandrasekaran, 2017). These models help researchers understand the underlying principles of sensory integration, such as reliability-based cue weighting and inverse effectiveness (Stein, Stanford, & Rowland, 2020). By comparing model predictions with empirical data, researchers can test hypotheses about the neural mechanisms of multisensory integration and gain insights into how the brain creates coherent perceptions from diverse sensory inputs (Blohm et al., 2020).

A wide range of computational models for multisensory integration is available (Colonius & Diederich, 2020), providing descriptions at computational, algorithmic, and implementation levels within information-processing theory (Marr, 2010). Nevertheless, it is unusual to find formal evaluations of multisensory integration using more than one model at the time to bridge across levels of analysis (Ursino, Cuppini, & Magosso, 2014).

Scikit-NeuroMSI was designed to meet three fundamental requirements in the computational study of multisensory integration:

1. **Modeling Standardization:** Researchers need to compare different theoretical approaches (Bayesian, neural network, maximum likelihood estimation, among others) using consistent analysis methods. Our framework provides a standardized interface for implementing and analyzing different types of models.
2. **Data Processing Pipeline:** The framework handles multidimensional data processing across:
  - Spatial dimensions (1D to 3D spatial coordinates)
  - Temporal sequences
  - Multiple sensory modalities (e.g., visual, auditory, touch)
3. **Analysis Tools:** We provide integrated tools for:
  - Parameter sweeping across model configurations
  - Result visualisation and export
  - Statistical analysis of model outputs

Furthermore, the software processes fundamental attributes of experimental inputs pertinent to multisensory research: spatial coordinates (e.g., degrees of visual angle and sound source location), temporal properties (encompassing stimulus onset, duration, and inter-stimulus intervals), stimulus intensity, and spatiotemporal reliability (random noise). Each input is validated to ensure compliance with the formatting standards, followed by a transformation to standardized internal representations that facilitate effective model processing. The software systematically applies type checking and data validation protocols to ensure the integrity of the inputs before proceeding with processing.

Consequently, the software requires a uniform output for each multisensory integration model. This output incorporates the activity values from all participating modalities, necessitating at least two unisensory modes along with one multisensory mode. There is an optional provision for output related to causal inference responses, as determined by the user. These prerequisites guided the technical design choices detailed in the next section.

**Code 1.**
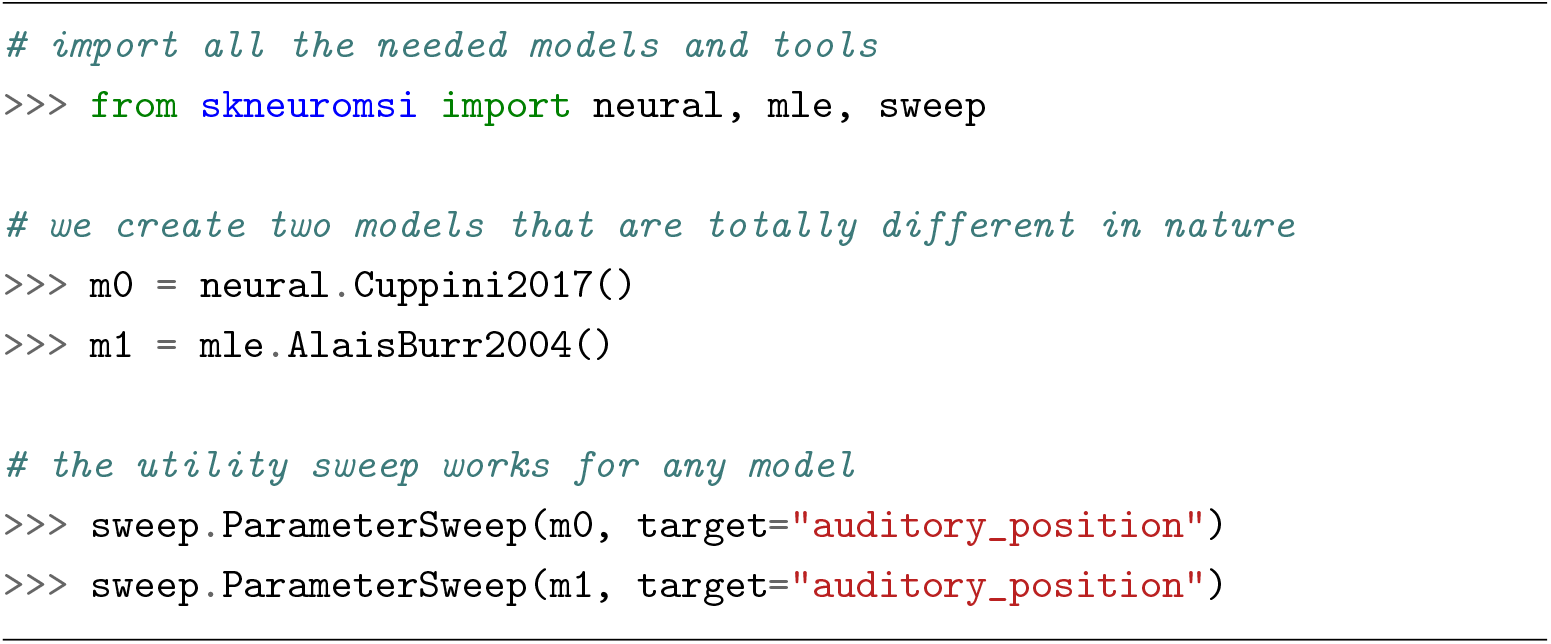
Example showing how Scikit-NeuroMSI enables the use of consistent analysis tools across different types of multisensory integration models, regardless of their underlying theoretical framework.

### 2.2 A formal and technical approach for the model standardization

We outline the theoretical and mathematical basis of multisensory integration models. We first describe the formal properties and mathematical framework needed to understand these computational problems theoretically, which aids in addressing software engineering challenges. Our development of `Scikit-NeuroMSI` aimed to articulate these problems with minimal conceptual differences, removing programming overhead to focus on solving multisensory integration issues (Brooks & Kugler, 1987). Specific implementation details and practical considerations are discussed in Section 2.3, with examples of applying these principles in the software framework.

#### 2.2.1 Computational model formalities

This work aims to create a unified framework for multisensory integration models (Colonius & Diederich, 2020) through Scikit-NeuroMSI, facilitating interoperability between analysis, comparison, and explanation tools, regardless of the specific model.

Consider two distinct model outputs *r*0 and *r*1, such as Bayesian models (Körding et al., 2007; Shams & Beierholm, 2010), neural networks (Cuppini, Magosso, Bolognini, Vallar, & Ursino, 2014; Cuppini et al., 2017), or maximum likelihood estimators (Alais & Burr, 2004; Ernst & Banks, 2002), with different modalities and dimensions. Any processing function *f* should handle both *r*0 and *r*1 equally well.

In general, our goal is to create models (*m*) that produce results (*r*) compatible with any processing function *f*, ensuring that the new *m* or *f* are mutually compatible (see Appendix A for a mathematical formulation). For example, Code 1 shows that neural network models (Cuppini et al., 2017) and maximum likelihood estimation models (Alais & Burr, 2004) can be analyzed with the same parameter sweep tools.

Another goal was to allow users to choose the sensory modality they simulate, reflecting real-world scenarios. For example, a researcher might study visual-auditory stimuli interaction in one experiment and visual-tactile in another. This allows models to adapt parameters to dynamically match selected modalities. As demonstrated in Code 2, an audio-visual setup reveals parameters such as `“auditory_position”`, while a visual-tactile setup replaces them with parameters such as `“tactile_position”`, keeping the core model structure and logic intact.

The final design requirement ensures multidimensional analysis of data. Each model output point is associated with a specific mode, time, and space, with the spatial dimension possibly encompassing one to three dimensions. Consequently, the model output can comprise up to five dimensions (5D).

#### 2.2.2 Object-oriented design

Selecting an ecosystem for a tool is a subjective decision. Most data analysis projects use Python due to its rich scientific ecosystem (Perez, Granger, & Hunter, 2010). We chose Python to exploit object-oriented mechanisms, enhancing simplicity and tool extensibility. In object-oriented languages such as Python, classes structure data types by grouping attributes (state) and methods (behavior), enabling cohesive code organization and providing inheritance for sharing functionalities without code duplication (Booch, 1982).

A conceptual mechanism available in object-oriented programming languages that we employed is abstract classes. These represent data types that, although possessing a complete protocol (i.e., their functions, the data they receive, and the return types are fully defined), have certain behaviors that remain unspecified. For example, as demonstrated in code 3, the `Foo` class utilizes the functionality already established in the `method0()` method without necessitating redundancy in the source code. Furthermore, this establishes a hierarchical structure, or inheritance, between the `FooABC` and `Foo` types, indicating that any object created or instantiated by `Foo` is an instance of the `FooABC` type. The final aspect to note is that `FooABC` is explicitly declared as an incomplete entity and is thus non-instantiable, achieved by adorning `method1()` with @abstractmethod.

We followed the “Dependency Inversion Principle” (DIP), an essential SOLID principle (R.C. Martin, 2000), which establishes two fundamental rules:

1. High-level modules should not depend on low-level modules. Both should depend on abstractions.
2. Abstractions should not depend on details. Details should depend on abstractions.

To illustrate its importance, we compare two model processing tool implementations. Code 4 is tightly coupled, relying directly on a specific model, while Code 5 is more flexible, using dependency injection via an abstract base class.

The main distinction between these methodologies lies in that Code 4 contains a tool function specifically hardcoded to operate solely with `Model1`, thereby precluding its application to other model types unless the function is modified. In contrast, Code 5 is designed to accommodate any model inheriting from `ModelABC`, facilitating the seamless integration of new model types, allowing runtime model substitution, enhancing testability through mock objects, and ensuring a more robust separation of concerns. This pattern is essential for Scikit-NeuroMSI, as it enables the consistent implementation of tools across various multisensory integration models.

**Code 2.**
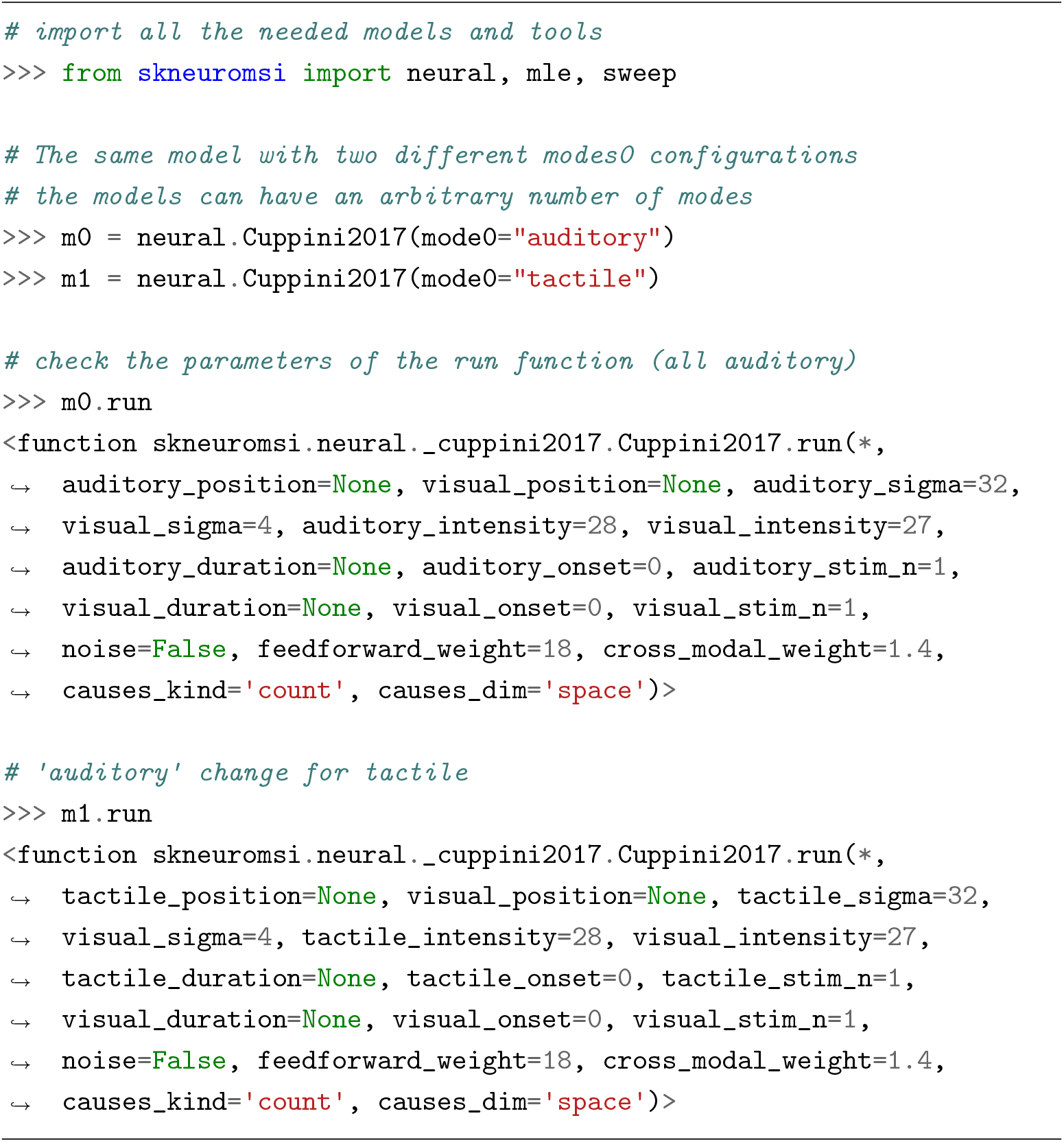
Example of mode configuration in a model: The same neural network model (`Cuppini2017`) can be configured for different sensory modalities. Compare how the run function parameters automatically adapt between auditory-visual (top) and tactile-visual (bottom) configurations while maintaining the same underlying model architecture. Note that the outputs of `m0.run` and `m1.run` show the function signatures since they are not executed. To execute the function, the code should be `m0.run()` or `m1.run()`

**Code 3.**
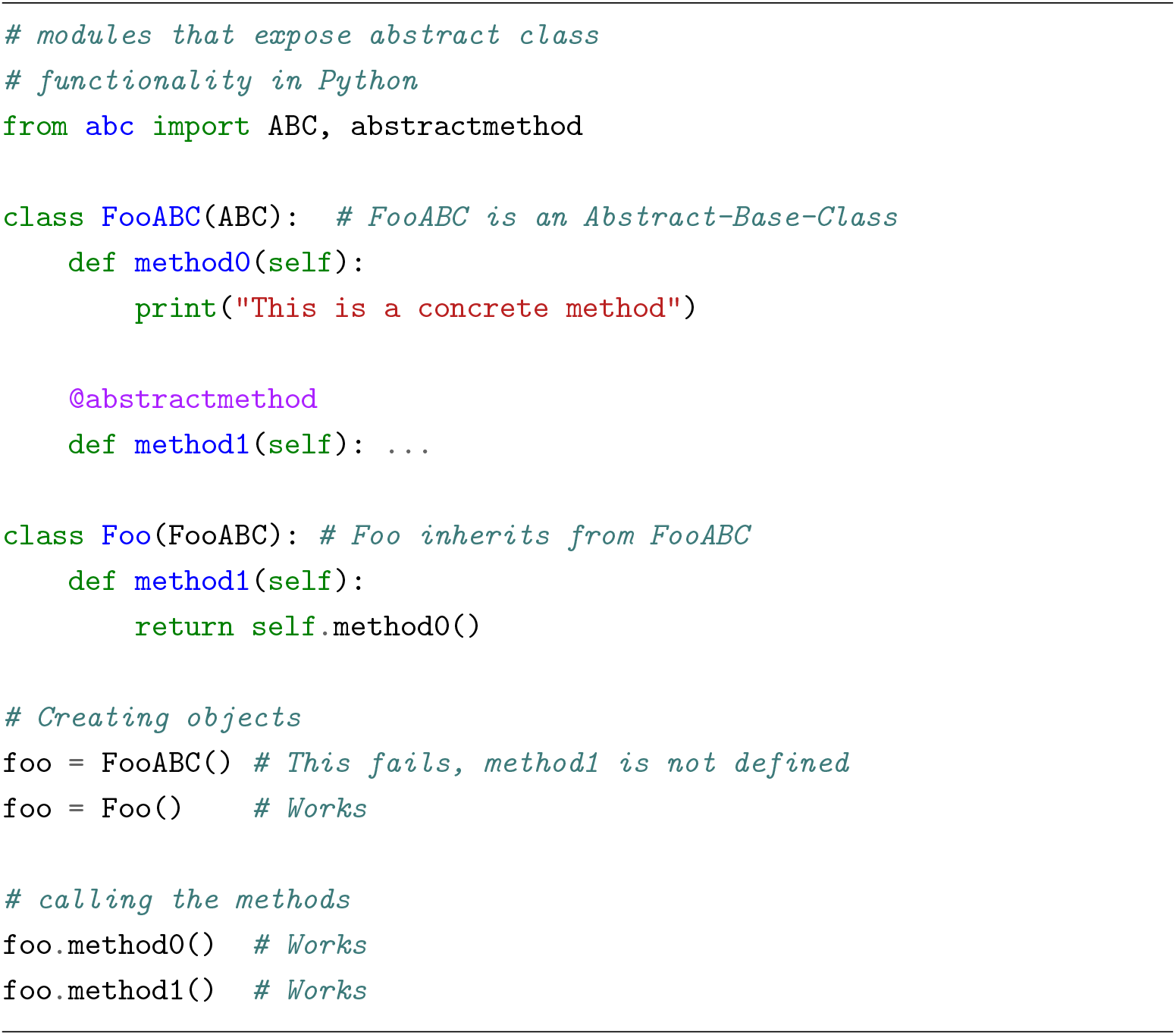
Example of abstract vs concrete class implementation in Python: The abstract class (`FooABC`) defines a protocol with both concrete and abstract methods, while its concrete implementation (`Foo`) provides the required implementation of the abstract method. Note how creating an instance of the abstract class fails, while the concrete class is instantiated successfully.

We propose a set of classes to reduce the “semantic gap” that separates the ideas of how a multisensory integration problem is expressed and how the code that represents these ideas is written. Two main classes form the core architecture of the framework:

- `ModelABC`: An abstract base class that defines the standard interface for all multisensory integration models
- `NDResult`: A result object responsible for storing multidimensional stimulus information and providing analysis tools for research

**Code 4.**
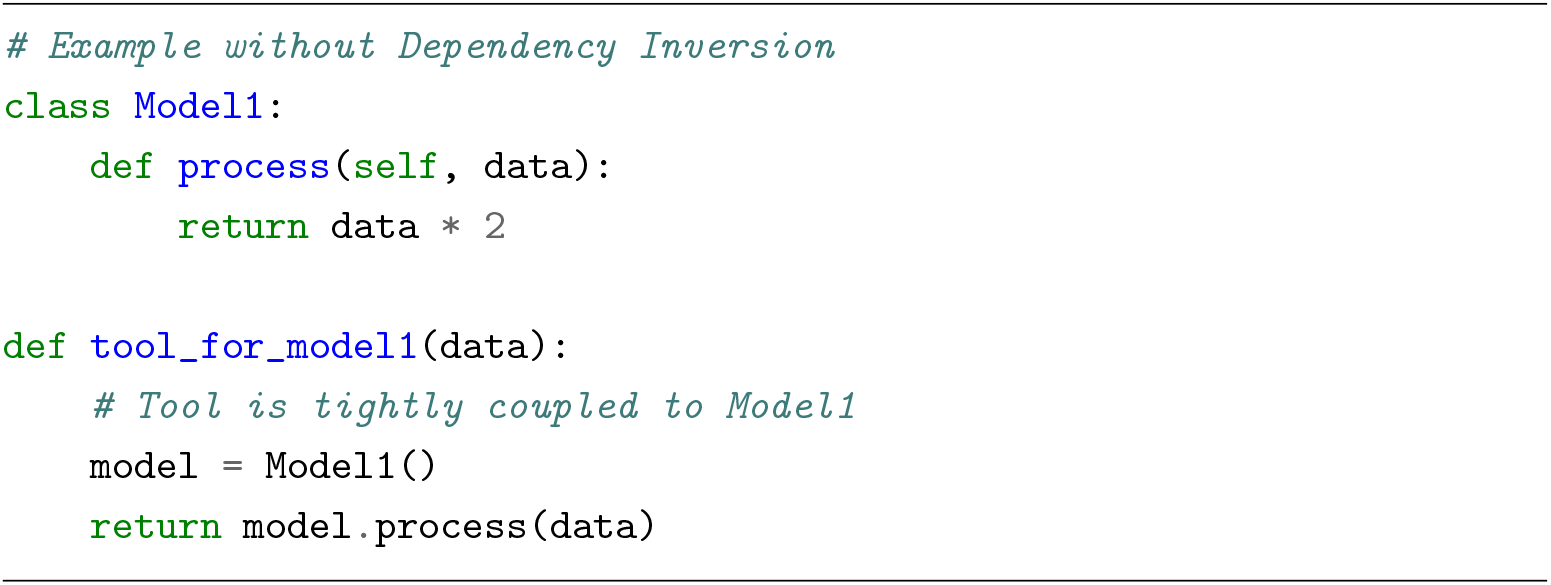
Example of a tool without Dependency Injection: The tool function is tightly coupled to a specific model implementation (`Model1`), making it inflexible and difficult to extend to other model types.

**Code 5.**
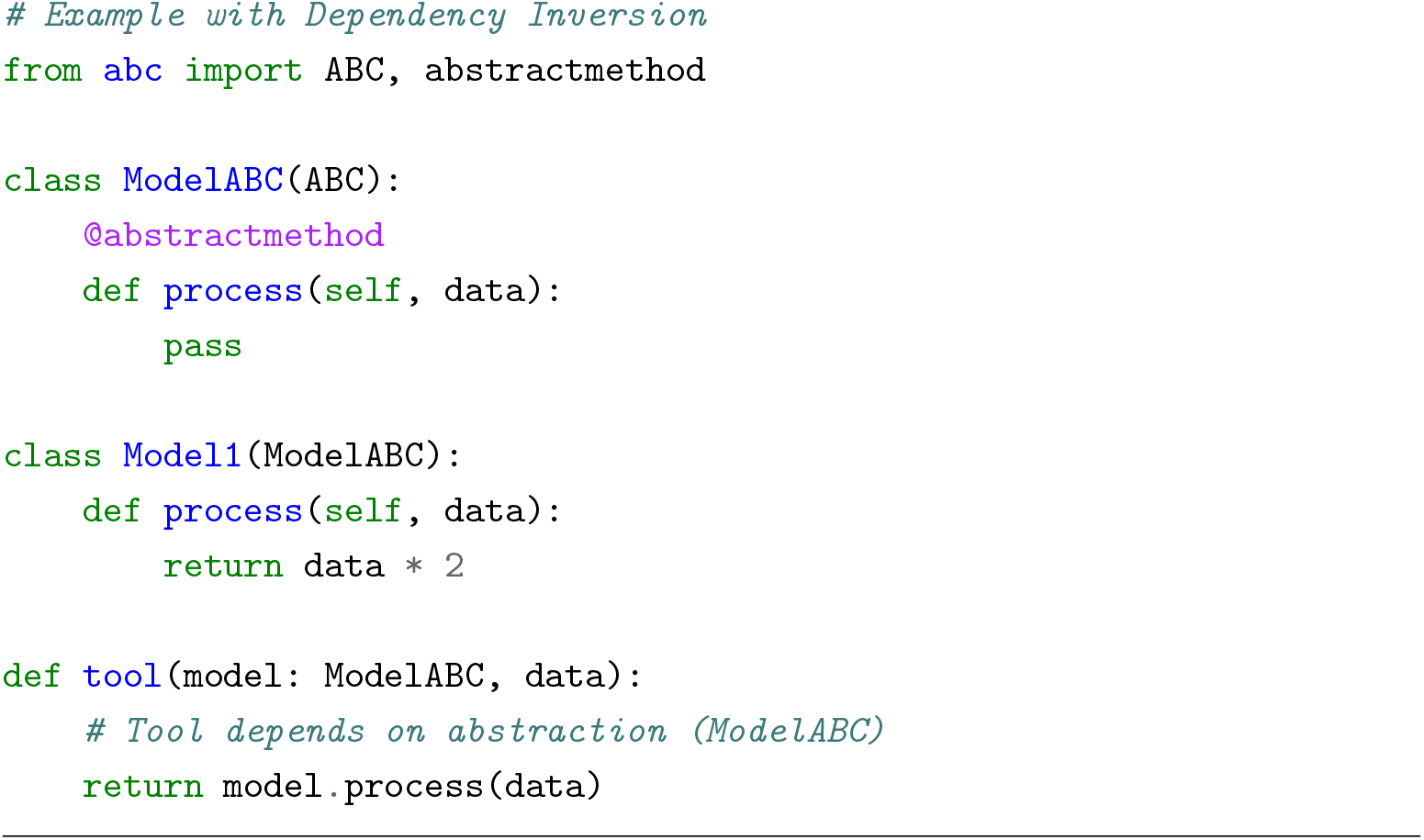
Example of a tool with Dependency Injection: Depending on an abstraction (`ModelABC`), the tool function can work with any model that implements the required interface, allowing flexible model substitution and easier testing.

This two-class design separates the model implementation logic from the data handling and analysis capabilities, following standard software engineering principles. The entirety of the classes, along with their interconnections, is presented in a diagram employing UML language (Jacobson, Booch, & Rumbaugh, 2000) to formally depict the relationships among the core modules of the project (see Figure 1).

**Fig. 1.**
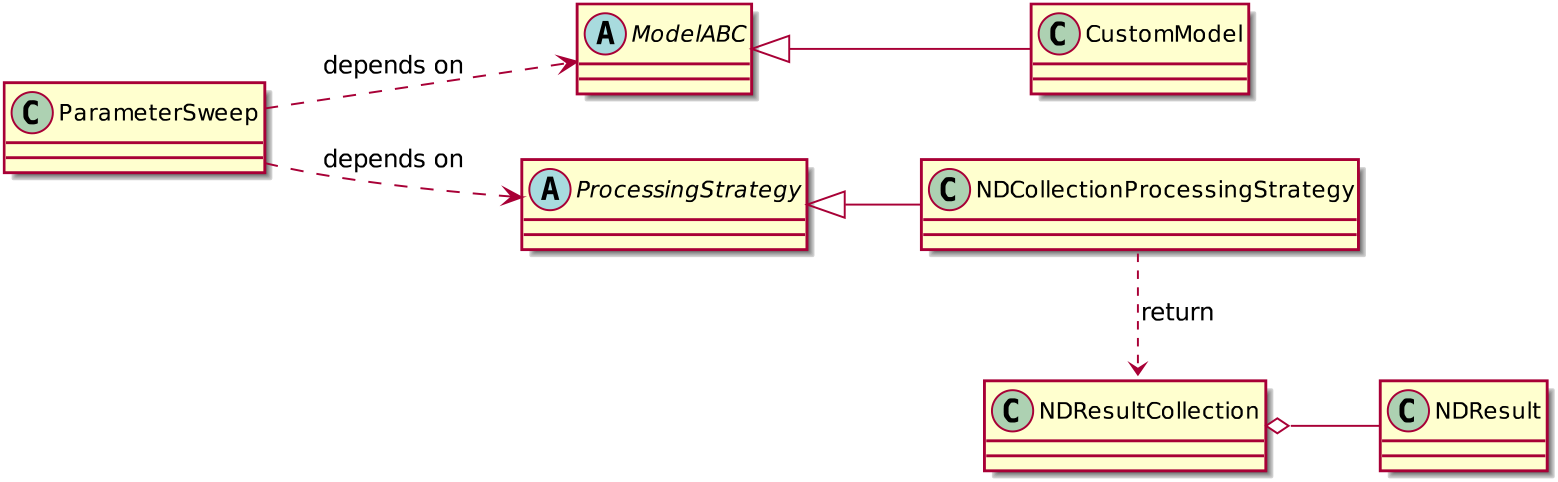
Reduced Class Diagram of *Scikit-NeuroMSI*. Empty arrows represent inheritance, the diamond-headed arrow indicates that multiple `NDResult` objects are aggregated into a single `NDResultCollection`, and all other relationships have explanatory labels

ParameterSweep is a tool for performing parameter sweeps over models, offering high flexibility and employing interchangeable strategies to process results, thereby enabling efficient memory management. NDResultCollection serves as an auxiliary class designed to aggregate and compress results into organized collections. This facilitates advanced functionalities, such as the analysis of spatiotemporal disparity effects. Furthermore, NDResultCollection is the standard output format of ParameterSweep when using the default Processing Strategy. For a complete description of the implemented classes, refer to Appendix B.

### 2.3 Implemented Models

Any model of multisensory integration must define the link between responses to unisensory signals, such as visual and auditory, and responses to cross-modal signals such as visual-auditory. This connection varies according to spatial and temporal characteristics, experimental configuration, and level of description (e.g. single neurons, neural populations, neuroimaging, behaviors). This opens a broad spectrum of approaches for modeling the observations derived from multisensory integration experiments.

A prevalent paradigm in this field is the Ventriloquist Effect (Thurlow & Jack, 1973). This phenomenon arises when incongruent visual and auditory stimuli are simultaneously presented, leading the observer to perceive a singular origin for both visual (the movements of a puppet’s face) and auditory (speech) stimuli, attributing them to the same source (the puppet’s speaking) (see Figure 2 for an illustration). This effect is systematically investigated in laboratory environments through tasks designed to assess the spatial localization of an auditory source within a combined visual-auditory setting (Alais & Burr, 2004).

**Fig. 2.**
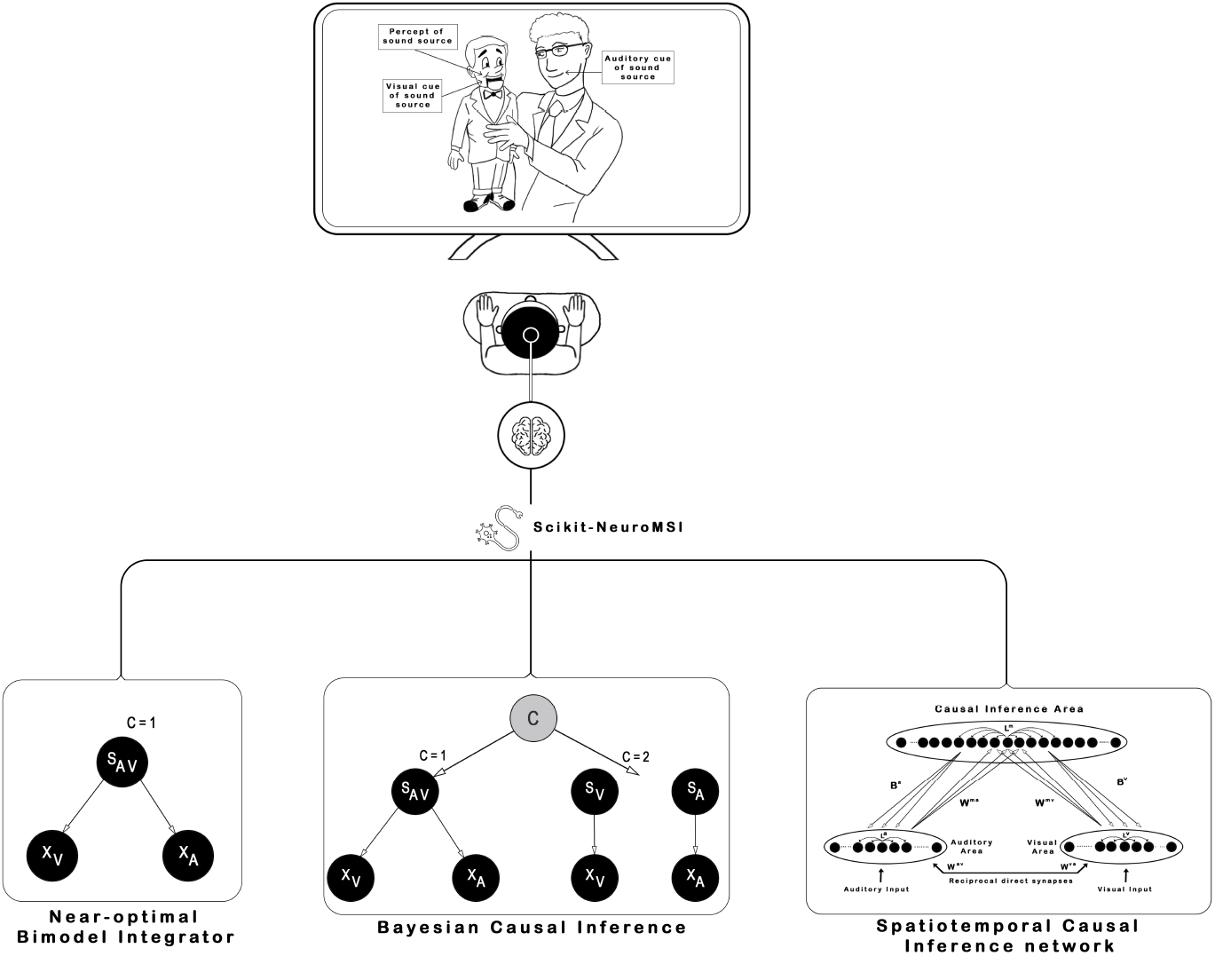
Implemented models in *Scikit-NeuroMSI*. The illustration represents how the software package allows to model the Ventriloquist Effect (i.e. spatial integration under audio-visual disparities) using three different approaches: Near-optimal Bimodal Integrator, Bayesian Causal Inference and Spatiotemporal Causal Inference network

In general, these models aim to explain common “empirical rules” derived from multisensory integration paradigms (see Colonius and Diederich (2020) and Stein et al. (2020) for a detailed review). The “spatio-temporal rule” suggests that stimuli near in space and time are more likely to be integrated. The “inverse-effectiveness rule” states that integration is stronger when the unimodal input intensity decreases. Additionally, the “reliability rule” emphasizes a greater weighting of more reliable modalities (e.g., with lower noise). These empirically observed “rules” have been demonstrated to arise from common computational principles (Ohshiro et al., 2011) and do not necessarily require alternative models for their explanation. In consequence, the key challenge is the development of models capable of accounting for empirical observations of multisensory integration across various levels of description.

Here we present the main models of the Ventriloquist Effect currently available in the Scikit-NeuroMSI package. Appendix C contains a comprehensive mathematical exposition of the models, as well as the code necessary for their execution. The package currently includes models pertinent to additional paradigms, such as the Sound-Induced Flash Illusion (Cuppini et al., 2014; Zhu, Beierholm, & Shams, 2024b). These models are not elaborated upon in depth in this article, but are thoroughly delineated in the user documentation. Furthermore, we provide guidance on how to streamline the incorporation of novel models into the package. We actively encourage the research community specializing in multisensory integration to contribute to this initiative by developing and sharing their own models (refer to Contributing Guidelines).

#### 2.3.1 Near-optimal Bimodal Integrator

An early model proposes that the process of cue combination from different modalities resembles a maximum-likelihood integrator (Ernst & Banks, 2002). The Near-optimal Bimodal Integrator (Alais & Burr, 2004) for auditory (A) and visual (V) signals in the context of an auditory spatial localization task (e.g. Ventriloquist effect) can be computed by adding the unisensory estimates (*Ŝ*_*A*_ and *Ŝ*_*V*_) weighted by their reliability. Consequently, the integrated percept is predisposed to align more closely with the signal that exhibits lower variability.

#### 2.3.2 Bayesian Causal Inference

In the previous model, cue integration is essential for accurately estimating a cross-modal stimulus. However, if there is a significant difference between the subjective assessments *Ŝ*_*A*_ and *Ŝ*_*V*_ from a visual-auditory stimulus, the observer cannot determine if this difference is due to random noise in neural signal processing or systematic signal divergence.

Originally designed to explain the Ventriloquist Effect, the Bayesian Causal Inference model (Körding et al., 2007) distinguishes whether *Ŝ*_*A*_ and *Ŝ*_*V*_ arise from a single audiovisual event (integration) or separate events (segregation). To do so, the observer considers the likelihood of *Ŝ*_*A*_ and *Ŝ*_*V*_ given a common or separate event and the prior probability of a common source. A higher likelihood occurs if the two unisensory signals are similar, which in turn increases the probability of inferring that the signals have a common cause.

#### 2.3.3 Network model for audio-visual integration and causal inference

Bayesian causal inference models provide a high-level description (i.e. computational level of analysis according to Marr (Marr, 2010)) of the computations carried out by the brain to integrate unisensory signals. Recently, neural network models have been proposed as an alternative to model causal inference in multisensory integration paradigms (Cuppini et al., 2017; Fang, Yu, Liu, & Chen, 2019; Rideaux, Storrs, Maiello, & Welchman, 2021), providing a low-level description of such a mechanism.

The audio-visual integration and causal inference network (Cuppini et al., 2017) features two unisensory regions for processing noisy auditory and visual stimuli, inter-connected by cross-modal excitatory synapses. Here, rate-coded neurons are spatially organized, with closer neurons responding to nearer spatial positions. These regions emulate sensory processing in the brain’s unisensory cortex and determine the spatial location of the stimuli by computing the barycenter of activity in the auditory and visual regions. Cross-modal connections cause the spatial localization of one modality to be influenced by the concurrent presentation in another, even if processed separately.

#### 2.3.4 Multisensory Spatiotemporal Causal Inference Network

Our research group developed the Multisensory Spatiotemporal Causal Inference Network to account for the sound-induced flash illusion (Paredes, Ferri, Romei, & Seriès, 2025). This model was built upon preceding network models for spatial (Cuppini et al., 2017) and temporal (Cuppini et al., 2014) multisensory integration to inform two levels of causal inference processing. Our model consists of three layers: two encode auditory and visual stimuli separately and connect to a multisensory layer via feedforward and feedback synapses. At the unisensory areas, the model computes the spatiotemporal position of the external stimuli. In addition, at the multisensory area the model computes causal inference. This neural architecture allows iterative computation of spatiotemporal causal inference across the network (Rohe et al., 2019).

In summary, our model retains neural connectivity (lateral, cross-modal, feedforward) and inputs as detailed in the previously discussed network (Cuppini et al., 2017), while integrating feedback connectivity. Additionally, this model introduces latency into the cross-modal and feed-forward-feedback neural inputs in accordance with literature indicating that early and late interactions during multisensory processing. Furthermore, in line with Cuppini et al. (2014), temporal filters have been incorporated for auditory, visual, and multisensory neurons to replicate the temporal progression of neural input and synaptic dynamics. These filters account for specific time constants that determine the temporal characteristics of each group of neurons, namely auditory, visual, or multisensory.

## 3 Example Applications

### 3.1 Modeling Spatiotemporal Causal Inference with Scikit-NeuroMSI

Causal inference is a highly relevant computation for multisensory integration (Körding et al., 2007; Shams & Beierholm, 2022, 2010). Causal inference in multisensory integration is examined through implicit or explicit tasks. Explicit causal inference involves tasks in which participants are required to directly assess the causal relationship between stimuli (unity judgment) in a multisensory setting, whereas implicit causal inference involves tasks where participants are required to estimate the spatiotemporal location of the stimuli. An in-depth investigation of how the causal mechanism operates in both types of tasks is currently in progress (Acerbi, Dokka, Angelaki, & Ma, 2018) and has been found to be distinct in neurodiverse populations (e.g. Noel, Shivkumar, Dokka, Haefner, and Angelaki (2022)).

In general, multisensory causal inference models rely mainly on Bayesian inference (Körding et al., 2007), offering a high-level description (as described by Marr’s computational level (Marr, 2010)) of how the brain integrates sensory signals (French & DeAngelis, 2020). Recently, neural network models have emerged as an alternative for modeling causal inference in multisensory contexts (Cuppini et al., 2017; Fang et al., 2019; Rideaux et al., 2021), providing a tentative implementation of the mechanism. Yet, these models have not been rigorously tested across multiple experimental frameworks or dimensions (i.e. space and time), nor have they been compared with other models. In the following, we ask: 1) Are the probabilistic and network models comparable in their performance when fitting data? 2) Do these models accurately account for both implicit and explicit causal inference responses? 3) Do these models show performance differences when working with spatial or temporal disparities?

#### 3.1.1 Modeling Setup

We used the models currently implemented in Scikit-NeuroMSI to reproduce human responses in audio-visual causal inference tasks (see Experiments 2, 3 and 4 in Noel, Shivkumar, et al. (2022) for details on the experimental setup) and qualitatively compared their performance. For each task, we fitted the implemented models to behavioral responses of healthy control participants using the differential evolution algorithm (Storn & Price, 1997) available in the SciPy library for the Python programming language (Virtanen et al., 2020). Details about the fitting procedure and model readout for each task can be found in the Appendix D.

First, we model an auditory spatial localization task with disparate visual cues (Experiment 2 in Noel, Shivkumar, et al. (2022)) to examine the implicit causal inference performance of the models. Our main focus is the modulation of auditory spatial perception by visual stimuli, a phenomenon known as auditory bias. We present each model with an auditory stimuli at a fixed position (45°) and visual stimuli at different positions relative to the auditory cue: ±3, ±6, ±12, and ±24° (see Figure 3, left). For simplicity, we present each possible combination of audio-visual stimuli only once, and all noise sources are eliminated for this approach. Following Körding et al. (2007) and Cuppini et al. (2017), we compute the auditory bias as the spatial disparity between the position of the auditory stimulus and the position detected by each model, divided by the distance between the auditory and visual stimuli. We fit the auditory bias responses of the model to correspond with human responses under the high visual cue reliability condition (see Noel, Shivkumar, et al. (2022) for details).

**Fig. 3.**
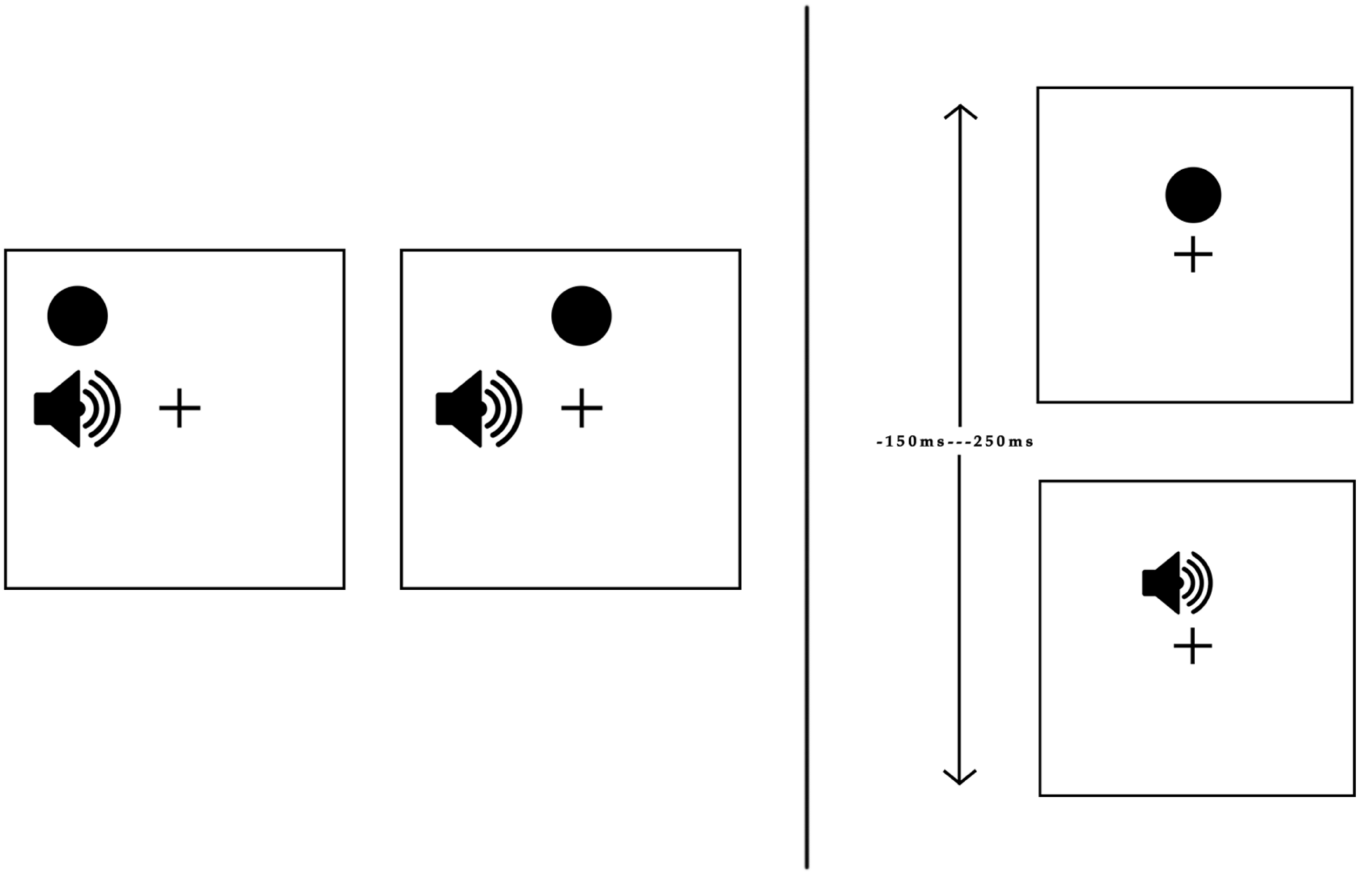
Causal inference tasks. The figure shows the causal inference tasks reported in Noel, Shivkumar, et al. (2022) that were simulated in this study. Left panel: Spatial audio-visual disparity task. Participants viewed a visual disk and heard an auditory tone at different locations and with different small disparities (top = no disparity, bottom = small disparity). We present each model with an auditory stimuli at a fixed position (45°) and visual stimuli at different positions relative to the auditory cue: ±3, ±6, ±12, and ±24°. The models had to determine the position of the auditory stimuli and report the number of causes (1 or 2). Right panel: Temporal audio-visual disparity task. Participants viewed a visual disk and heard an auditory stimuli at different onsets relative to the visual cue. We present each model with a visual stimulus at a fixed onset (160 ms) and auditory stimuli at different onsets relative to the visual cue: 0, ±20, ±80, ±150, and +250 ms. The models had to determine the number of causes of the stimuli (1 or 2)

Next, we model an audio-visual common cause report task under spatial disparities (Experiment 3 in Noel, Shivkumar, et al. (2022)) to examine the explicit causal inference performance of the models. Audio-visual stimuli are delivered to each model at the same positions as in the previous simulation. Here, our focus is the modulation of the unity judgments (common cause reports) of the models by the spatial disparity of the stimuli. For the Bayesian model, we determine the proportion of reports indicating a common cause by calculating the posterior probability of a common cause (refer to Equation C7). In contrast, for network models, this proportion is identified through the maximal neural activation (refer to Equation C9) observed within the multisensory neurons.

Finally, we model an audio-visual common cause report task under temporal disparities (Experiment 4 in Noel, Shivkumar, et al. (2022)) to examine the explicit temporal causal inference performance of the models. Here our focus is on how temporal disparities in stimuli affect the unity judgments of the models. We present each model with a visual stimulus at a fixed onset (160 ms) and auditory stimuli at different onsets relative to the visual cue: 0, ±20, ±80, ±150, and +250 ms (see Figure 3, right). Each possible combination of audio-visual stimuli is presented only once, and all noise sources are eliminated for this approach. We calculate the proportions of the common cause reports of the models as in the previous simulation.

#### 3.1.2 Simulation Results

We compared the performance of the implemented models in the aforementioned causal inference tasks. Auditory bias responses are shown in Figure 4a. We observe that both Bayesian (Körding et al., 2007) and Network models (Cuppini et al., 2017) provide a good approximation to behavioral data, while the maximum likelihood estimation model (MLE) (Alais & Burr, 2004) fails to reproduce audio-visual disparities beyond ±6°.

**Fig. 4.**
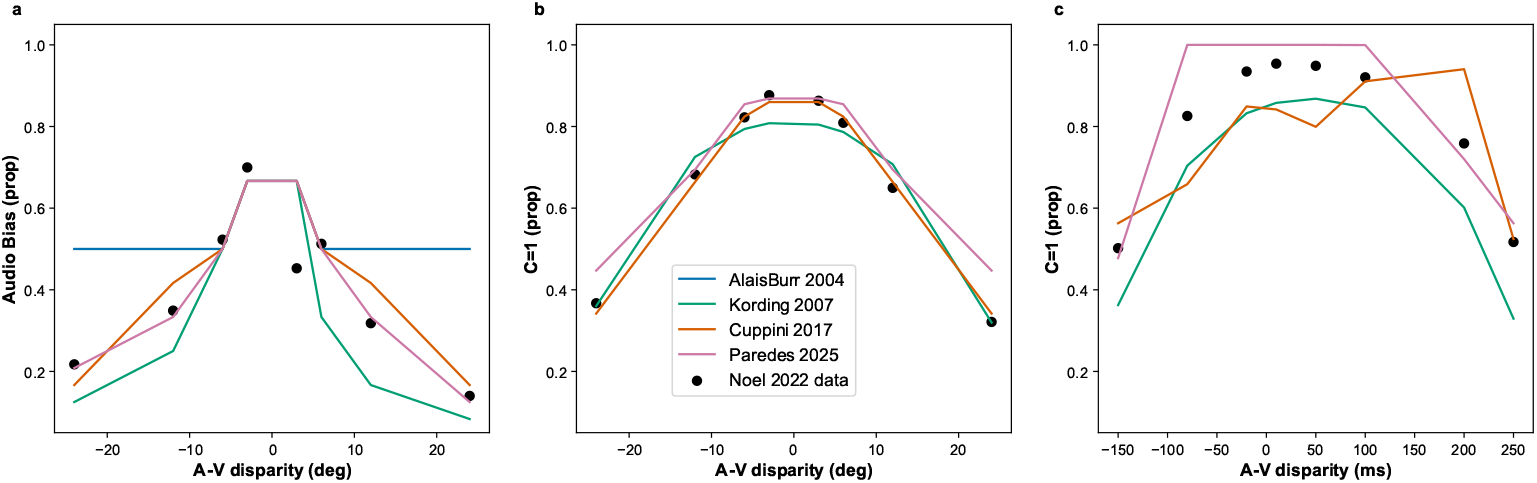
Models of audio-visual causal inference tasks. The figure shows optimal responses of the implemented models in multisensory causal inference in the audio-visual disparity tasks reported in Noel, Shivkumar, et al. (2022). **(a)** Performance of the models in spatial localization within an implicit causal inference task. This graph reveals that Bayesian and network models provide optimal performance, whereas the MLE model fails to reproduce audio-visual disparities beyond ±6°. **(b)** Performance of the models in common source reports under spatial disparities within an explicit causal inference task. These simulations show that the network models outperform the Bayesian Causal Inference model in audio-visual spatial disparities below ±6°. **(c)** Performance of the models in common source reports under temporal disparities within an explicit causal inference task. This graph shows that both the Multisensory Spatiotemporal Causal Inference Network and the Bayesian Causal Inference models provide fair approximations to human performance, whereas the network model for audio-visual integration fails to reproduce disparities beyond ±100 ms

Furthermore, spatial causal inference responses are shown in Figure 4b. We observe that all the evaluated models provide a fair approximation to behavioral responses, with the neural network models (Cuppini et al., 2017) outperforming the Bayesian Causal Inference model (Körding et al., 2007) in audio-visual disparities below ±6°.

Temporal causal inference responses are shown in Figure 4c. We observe that both the Bayesian Causal Inference model (Körding et al., 2007) and the Multisensory Spatiotemporal Causal Inference Network (Paredes et al., 2025) provide a good approximation to behavioral responses, whereas the network model for audio-visual integration (Cuppini et al., 2017) fails to reproduce temporal disparities beyond ±100 ms.

The findings from the current series of simulations indicate that the Bayesian Causal Inference and Spatiotemporal Causal Inference network models offer the most accurate representation of the participants’ data. However, the efficacy of both models diminishes when addressing temporal disparities, underscoring the need for new models that effectively incorporate causal inference within the temporal domain.

### 3.2 Comparing network and Bayesian models of Spatiotemporal Causal Inference with Scikit-NeuroMSI

To our knowledge, this is the first time that implicit and explicit spatiotemporal causal inference is computationally modeled using a combined Bayesian and Neural Network approach (Acerbi et al., 2018; Ursino et al., 2014). This methodology facilitates the concurrent assessment of the influence of each model parameter and the identification of similarities among them. We are now equipped to conduct comprehensive sweeps of parameters within each model and assess their effects on model responses to discern commonalities that may allow us to bridge both levels of description. This process enables the exploration of interindividual variability by combining different modeling approaches, offering new ways to study multisensory integration differences seen in psychiatric or neurological conditions (Cascio et al., 2012; Festa et al., 2017; Hahn et al., 2014; Haß et al., 2017; B. Martin et al., 2013; Noel, Paredes, et al., 2022; Paredes et al., 2022; Ramkhalawansingh et al., 2017; Stevenson et al., 2014; Wu et al., 2012; Zhou et al., 2018; Zvyagintsev et al., 2017). In the following, we ask: 1) What would the correlates of the components of Bayesian models be in a more neural implementation? 2) Do the parameters of each model have the same impact on implicit and explicit causal inference responses? 3) Are the effects of model parameters the same for tasks involving spatial or temporal disparities?

#### 3.2.1 Modeling Setup

For the Bayesian Causal Inference model, we explored the impact of varying the prior probability of a common cause (*p*_*common*_) and the precision of the unisensory estimates (*σ*_*a*_ or *σ*_*v*_). For Spatiotemporal Causal Inference network model, we explored the impact of manipulating the weights of cross-modal 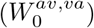, feedforward 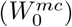, feedback 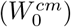 and excitatory lateral 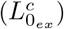 synapses. These parameters were selected due to its relevance in explaining individual differences in multisensory integration found in psychiatric conditions, as shown by recent computational research (Chrysaitis & Seriès, 2023; Karvelis, Seitz, Lawrie, & Seriès, 2018; Noel & Angelaki, 2023; Noel, Paredes, et al., 2022; Noel, Shivkumar, et al., 2022; Paredes et al., 2022).

First, we explored the impact of manipulating these parameters on the auditory bias responses of the selected models in the implicit causal inference task (Experiment 2 in Noel, Shivkumar, et al. (2022)). For simplicity, we computed the auditory bias at the −6° disparity point for each value of the explored parameters. Next, we examined the effects of sweeping parameters on the proportion of synchronous reports across spatial and temporal disparities (Experiments 3 and 4 in Noel, Shivkumar, et al. (2022)). Following Noel, Shivkumar, et al. (2022), these differences were systematically quantified by fitting Gaussian functions to the proportion of common source responses as a function of audio-visual disparities (Δ). The Gaussian fits provide three parameters that characterize the responses of the computational models: (1) amplitude, denoting the maximum proportion of common source reports by the model; (2) mean, indicating the Δ at which the proportion of common source reports was maximal; and (3) width (standard deviation), reflecting the extent of Δ within which the model was prone to report a common source.

#### 3.2.2 Simulation Results

The simulated auditory bias responses are shown in Figure 5a. We found that both *p*_*common*_ and *σ*_*v*_ within the Bayesian framework are inversely related to 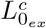 and 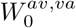 within the network level. In line with previous network modeling of audiovisual integration (Ursino, Crisafulli, di Pellegrino, Magosso, & Cuppini, 2017; Ursino et al., 2019), our results suggest a possible neural correlate of the prior probability of the co-occurrence of audio-visual stimuli in the cross-modal synapses, with such neural mechanism impacting unisensory precision as well.

**Fig. 5.**
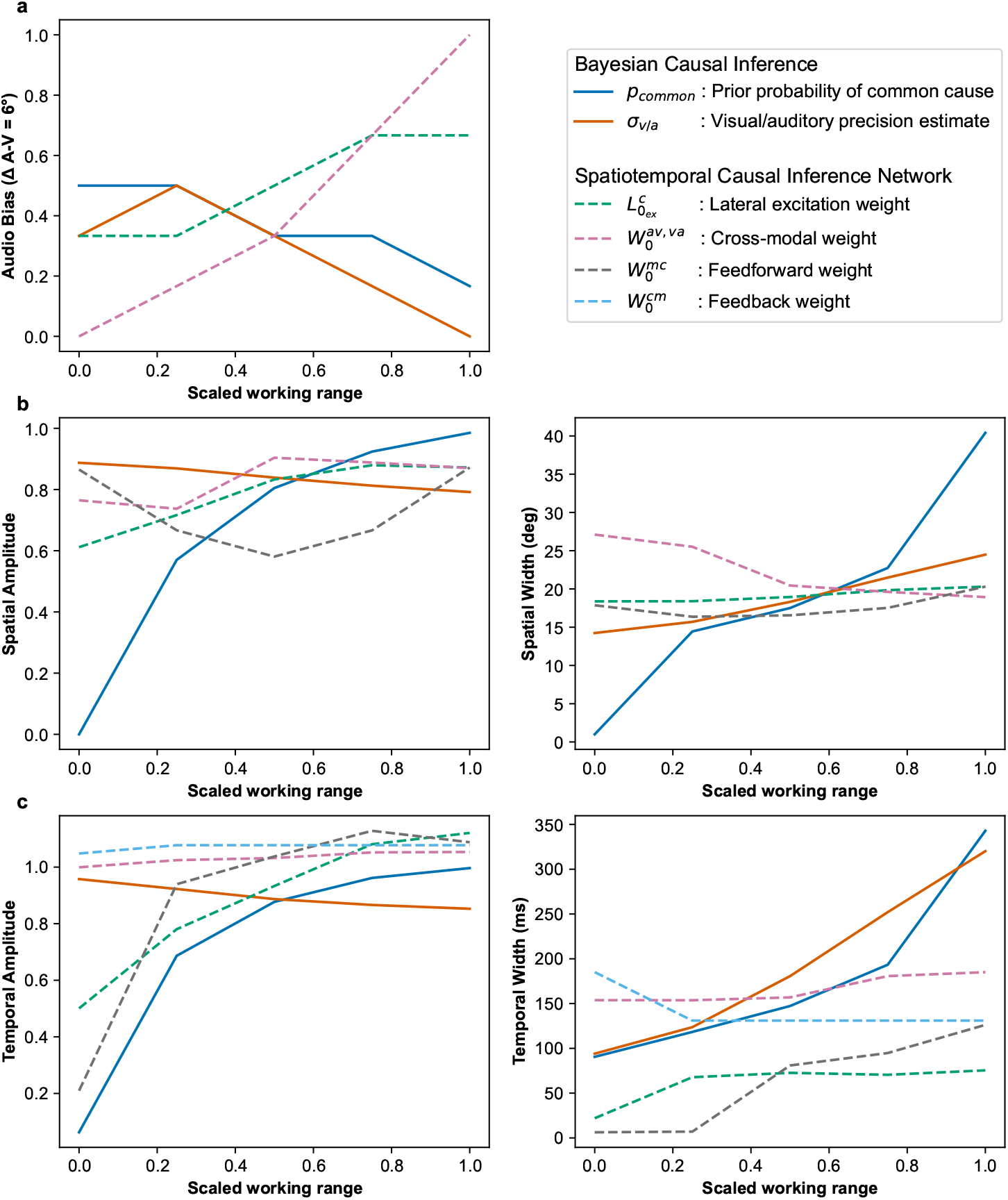
Impact of model parameters on causal inference responses in Bayesian and network models. **(a)** Parameter sweeps on the implicit causal inference task. The simulations indicate that the prior probability of a common cause (*p*_*common*_) and visual estimate precision (*σ*_*v*_) reduce auditory bias in the Bayesian model, while lateral excitation 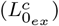 and cross-modal weights 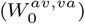 enhance it in our network model. **(b)** Parameter sweeps on the explicit spatial causal inference task. The proportion of common source responses relative to spatial disparities fitted to a Gaussian function for analysis. The simulations show that in the Bayesian Causal Inference model the parameter *σ*_*v*_ shows an opposite impact in the amplitude and width of the common source reports compared to 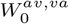 in the model. **(c)** Parameter sweeps on the explicit temporal causal inference task. The simulations show that in the Bayesian model *p*_*common*_ displays a similar impact in the amplitude and width of the common source reports compared to 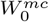 in the network model

In addition, the simulated common source responses in the spatial causal inference task are shown in Figure 5b. We found that *σ*_*v*_ within the Bayesian model shows an opposite impact in the amplitude and width of the common source reports compared to 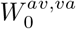 at the network level. This highlights the impact of cross-modal connectivity in explicit causal inference judgments, suggesting that its observed association with sensory precision estimates potentially scale up towards higher order cortical areas responsible for causal inference computations (Rohe et al., 2019; Rohe & Noppeney, 2015).

The simulated common source responses in the temporal causal inference task are shown in Figure 5c. In contrast to observations in the spatial domain, we found that the *p*_*common*_ within the Bayesian model displays a similar impact in the amplitude and width of the common source reports compared to 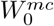 at the network level. This discrepancy opens up questions about potential differences in the mechanisms driving temporal and spatial causal inference, or at the very least, in the foundational assumptions under which these models were initially formulated. Notably, most of the modeling efforts have been carried out in spatial (static) multisensory integration tasks, whereas models of causal inference in the temporal domain at different levels of description have recently begun to accumulate (Cuppini et al., 2014; Pesnot Lerousseau, Parise, Ernst, & Van Wassenhove, 2022; Zhu et al., 2024b).

Overall, the Bayesian parameter *p*_*common*_ representing prior beliefs about common causes could be mapped to neural parameters such as 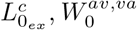 and 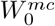 representing inter and intra-areal synaptic strengths, although no exact parallel could be found across domains (spatial or temporal) or metrics (bias, amplitude and width). Similarly, *σ*_*v/a*_ representing uncertainty in sensory information could be inversely mapped to 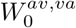, reflecting the strength of cross-modal connectivity at the network level, due to their opposite impact in the amplitude and width of the common source reports during explicit tasks. The observed discrepancies, including the differing effects of Bayesian and network models on implicit versus explicit causal inference tasks, indicate the presence of additional neural complexities that may not be fully encapsulated by Bayesian modeling or, conversely, by network approaches. The acquired understanding of the similarities of these models opens up the possibility of extending the current theoretical accounts of multisensory integration (Colonius & Diederich, 2020).

## 4 Discussion

We have addressed the objective of developing scientific software specifically designed for the computational modeling of multisensory integration, attending a key necessity in the field (Colonius & Diederich, 2020; Shams & Beierholm, 2022). We demonstrated the capabilities of Scikit-NeuroMSI in facilitating the implementation of multisensory integration models and systematically investigating their behavior by sweeping parameters across simulations (see Code 1 and Figure4). We have also demonstrated the utility of the software by modeling spatiotemporal causal inference at different levels of analysis using Bayesian (Körding et al., 2007) and network models of multisensory integration (see Figure 5), addressing a fundamental inquiry necessary for advancing the field (French & DeAngelis, 2020; Shams & Beierholm, 2022; Ursino et al., 2014).

With software tools such as a Scikit-NeuroMSI we are now able to approximate multisensory integration at different levels of analysis (Marr, 2010) (e.g. computational, algorithmic, and neural) simultaneously and extend our possibilities of generating computationally informed hypotheses. This enables the formulation of more precise predictions that can be evaluated with neurobiological and behavioral measurements, a factor crucial for the consolidation of emerging theories of multisensory integration in neuroscience (Colonius & Diederich, 2020; Shams & Beierholm, 2022). An immediate application for our new modeling framework is the study of multisensory integration differences in psychiatric and neurological disorders (Cascio et al., 2012; Festa et al., 2017; Hahn et al., 2014; Haß et al., 2017; B. Martin et al., 2013; Noel, Paredes, et al., 2022; Paredes et al., 2022; Ramkhalawansingh et al., 2017; Stevenson et al., 2014; Wu et al., 2012; Zhou et al., 2018; Zvyagintsev et al., 2017). For example, a novel quantitative theory on ASD (Noel & Angelaki, 2023) suggests that ASD could be interpreted as a multisensory causal inference disorder (computational level), where this process may be facilitated by divisive normalization (algorithmic level) and potentially disrupted by excitatory/inhibitory imbalances (neural implementation level). However, there is as yet no formal evaluation of experimental data of this disorder using more than one multisensory integration model at the time to bridge across levels of analysis.

We acknowledge that our modeling effort represents a first step towards achieving a general solution for multisensory integration formalization. We have shown the capabilities of our software in the simulation of multiple models in a group of three similar tasks (Noel, Shivkumar, et al., 2022). However, our software framework requires the incorporation of model comparison and validation metrics to facilitate the critical assessment of each model implementation (Blohm et al., 2020; Wilson & Collins, 2019). Overall, we propose a software environment as a first approach to a generalized framework for multisensory integration, needed for the theoretical advancement of the field.

## 5 Availability and Future Directions

The entire source code is under a BSD 3-Clause License and available in a public repository: https://github.com/renatoparedes/scikit-neuromsi. Scikit-NeuroMSI is available for installation on the Python Package-Index (PyPI)^1^. User documentation is automatically generated from Scikit-NeuroMSI docstrings and published in the Read the Docs service^2^.

In Spanish, there is a phrase “*Con el diario del Lunes*” (literally, “With Monday’s newspaper”), which shares the same meaning as the English expression “Monday-morning quarterback” - indicating that something becomes obvious only after it has happened. While Scikit-NeuroMSI has successfully achieved its technical objective of standardizing existing multisensory integration models, our experience has revealed opportunities for improved computational modeling.

Specifically, the architecture could be enhanced by decomposing the models into two distinct entities:

- A stimulus processing component that handles individual sensory inputs
- An integration component that consolidates the results into a unified modality

Or mathematically speaking:

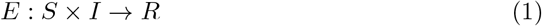

Where:

- *S* represents the stimulus source(s) from one or multiple modalities
- *I* is the integrator that combines the stimuli from *S*

We envision a Python implementation similar to what is presented in Code 6. By decoupling stimulus processing/generation from integration mechanisms, new integration models can be easily implemented and tested without modifying the underlying stimulus code. The flexible architecture simplifies the implementation of sophisticated integration strategies and enables straightforward extension to handle additional modalities or stimulus types, making the framework particularly valuable for emerging research in areas such as brain-computer interfaces and robotics. From a software perspective, this new design would promote code reusability and make the codebase more maintainable, allowing the scientific community to contribute new models and extensions to the framework more easily.

**Code 6.**
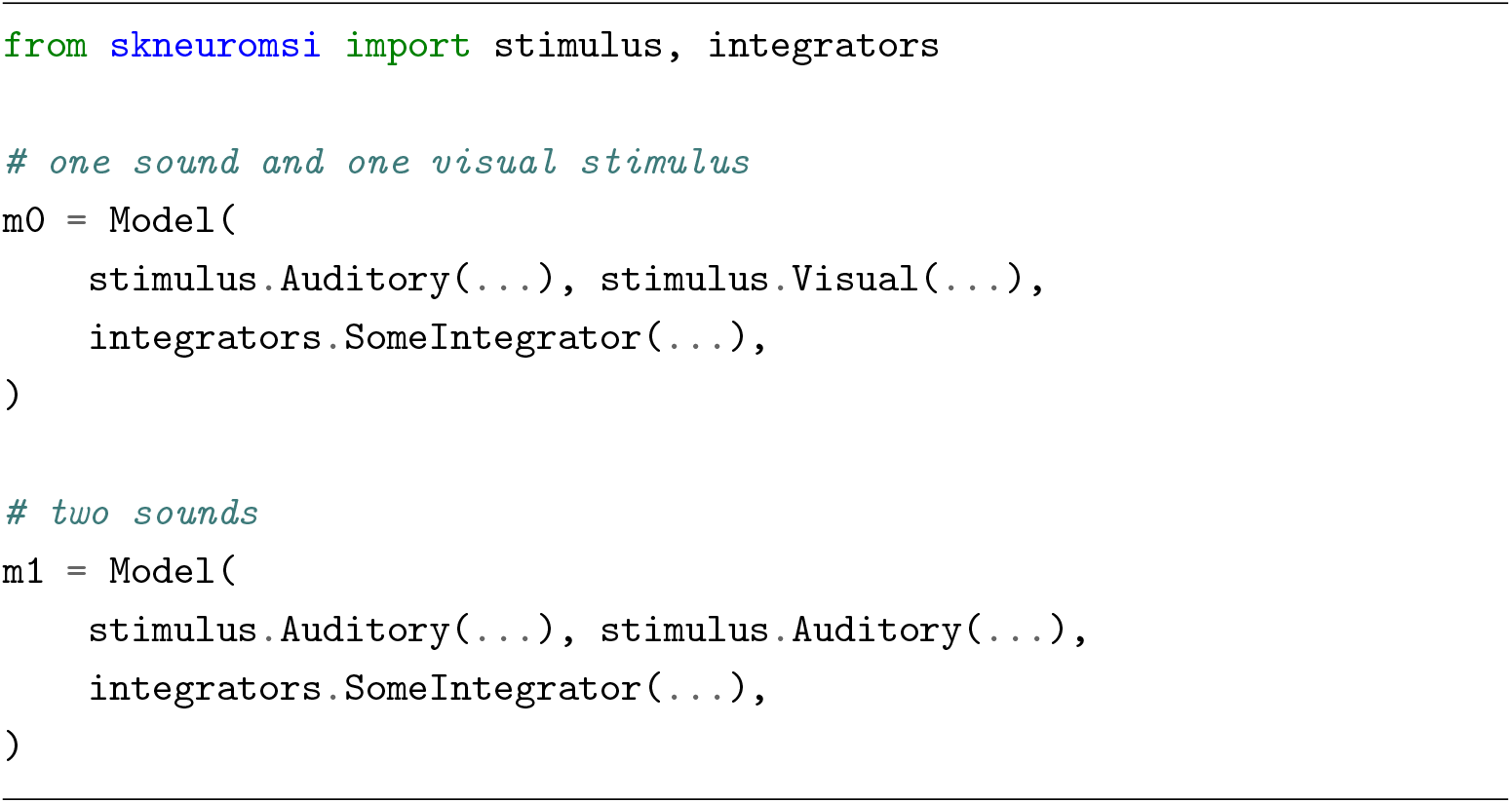
Proposed implementation using composition-based design: The same integration component (`integrators.SomeIntegrator`) can process both cross-modal (audio-visual) and unimodal (audio-audio) combinations through a flexible stimulus composition interface. This design separates stimulus handling from integration logic, enabling more modular and extensible implementations.

## Statements

### Authors Contribution

RP: Conceptualization, Methodology, Software, Visualization, Formal Analysis, Writing - original draft. JBC: Conceptualization, Methodology, Software, Visualization, Writing - original draft. PS: Supervision, Writing - reviewing and editing.

## Acknowledgments

The authors sincerely thank Jean-Paul Noel for providing the behavioral data modeled in this study and for providing insightful feedback on the preparation of this manuscript. The authors also express gratitude to Francisco Morote for contributing the illustrations included in the manuscript.

## Declarations

### Ethical Approval

Not applicable.

### Availability of data and materials

The code and data used to generate the simulations presented in this article is available at: https://github.com/renatoparedes/NeuroMSI-Network

## Appendix A Formal model standardization

Formally, our aim is to define a framework with the following property:

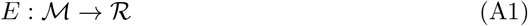

Where:

- *E* is the execution of a model *m*.
- ℳ is the set of all existing models.
- ℛ is the space of all possible results of ℳ.

Then for a specific model *m* ∈ ℳ

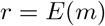

where *r* ∈ ℛ_*m*_ *⊆* ℛ.

Then we define *f* ∈ ℱ where ℱ represents all result processing tools, and we define a processing function:

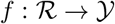

where 𝒴 is the output space of the tool; and with this we define the universal property

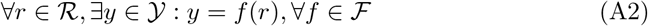

## Appendix B Classes and objects

The abstract class `ModelABC` defines the functionalities responsible for configuring mode names and establishing a common protocol for all models defined in the package. In terms defined in the previous equations, any model that inherits from `ModelABC` belongs to the set *M* of the equation A1.

The class `NDResult` is the implementation of any result from models inherited from `ModelABC` and from a mathematical point of view, they are the code realization of the set *R* from equation A1. As a data structure, `NDResult` (N-Dimensional Result) is a thin layer that adds multisensory integration analysis capabilities on top of the `DataArray` structure from the *XArray* library (Hoyer & Hamman, 2017). *XArray* was originally designed to work with high-dimensional data from NetCDF format (Rew & Davis, 1990), commonly used in satellite data processing. Additionally, since *XArray* provides persistence facilities, a `.ndr` persistence format was defined, which compresses a *NetCDF* file along with its metadata in a ZIP container (PKWARE Inc., 2022). Furthermore, `NDResult` also supports compression, as model results typically consume significant memory resources.

The `ParameterSweep` class is the primary tool designed to perform parameter sweeps over a model, automatically executing model multiple times while systematically varying the value of a target parameter within a specified range. The class allows configuration of the number of repetitions for each parameter value, supports parallel execution using multicore architectures, and implements memory management to prevent overflow conditions. Notably, `ParameterSweep` is model-agnostic and can work with any model that inherits from `ModelABC`, making it highly flexible and reusable. This is the first use of DIP.

The second use case implements the Strategy Pattern (Gamma, Helm, Johnson, Vlissides, & Patterns, 1995), allowing interchangeable algorithms to process the results of each model execution. By default, `ParameterSweep()` employs a strategy called `NDCollectionProcessingStrategy`, which acts as a processing pipeline that first compresses each individual result and then aggregates them into a coherent data structure called `NDResultCollection`, facilitating subsequent analysis. This architectural decision was driven by the memory footprint of the results, which typically occupy hundreds of megabytes, making concurrent storage of multiple results computationally expensive. The design allows for the implementation of custom strategies that can selectively store specific data points from each result, thus optimising memory utilisation.

`ParameterSweep` supports any model that instantiates from a class inheriting from `ModelABC`, and knows how to process the `NDResult` objects it generates by delegating them to the chosen strategy. In other words, it is the code implementation of the property defined in Equation A2.

`NDResultCollection` is the final piece of the puzzle that makes up Scikit-NeuroMSI. It is an auxiliary class that allows grouping many compressed results from the same model into a single collection and extends them with classic area functionalities such as cross-modal bias and causal analysis (Körding et al., 2007). Additionally, it is the default data type returned by the `ParameterSweep()` tool when used with the default strategy. Similar to `NDResult`, it offers the capability to export each data point to NetCDF format in a ZIP compressed file with its metadata in a format we call `*.ndc`.

## Appendix C Implemented models

### C.1 Near-optimal Bimodal Integrator

The Near-optimal Bimodal Integrator (Alais & Burr, 2004) for auditory (A) and visual (V) signals in the context of an auditory spatial localization task can be computed as:

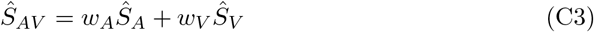

where *Ŝ*_*A*_ and *Ŝ*_*V*_ are unimodal auditory and visual estimates, respectively, and *Ŝ*_*AV*_ is the multimodal estimate.

In addition, *w*_*A*_ and *w*_*V*_ are the relative weights for each modality, defined as:

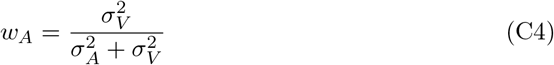

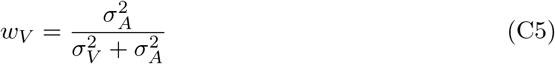

where *σ*_*A*_ and *σ*_*V*_ are the variances of each unimodal stimuli, respectively.

These equations show that the optimal multisensory estimate adds the unisensory estimates weighted by their normalized reciprocal variances.

The Near-optimal Bimodal Integrator model can be called using Code 7.

**Code 7.**
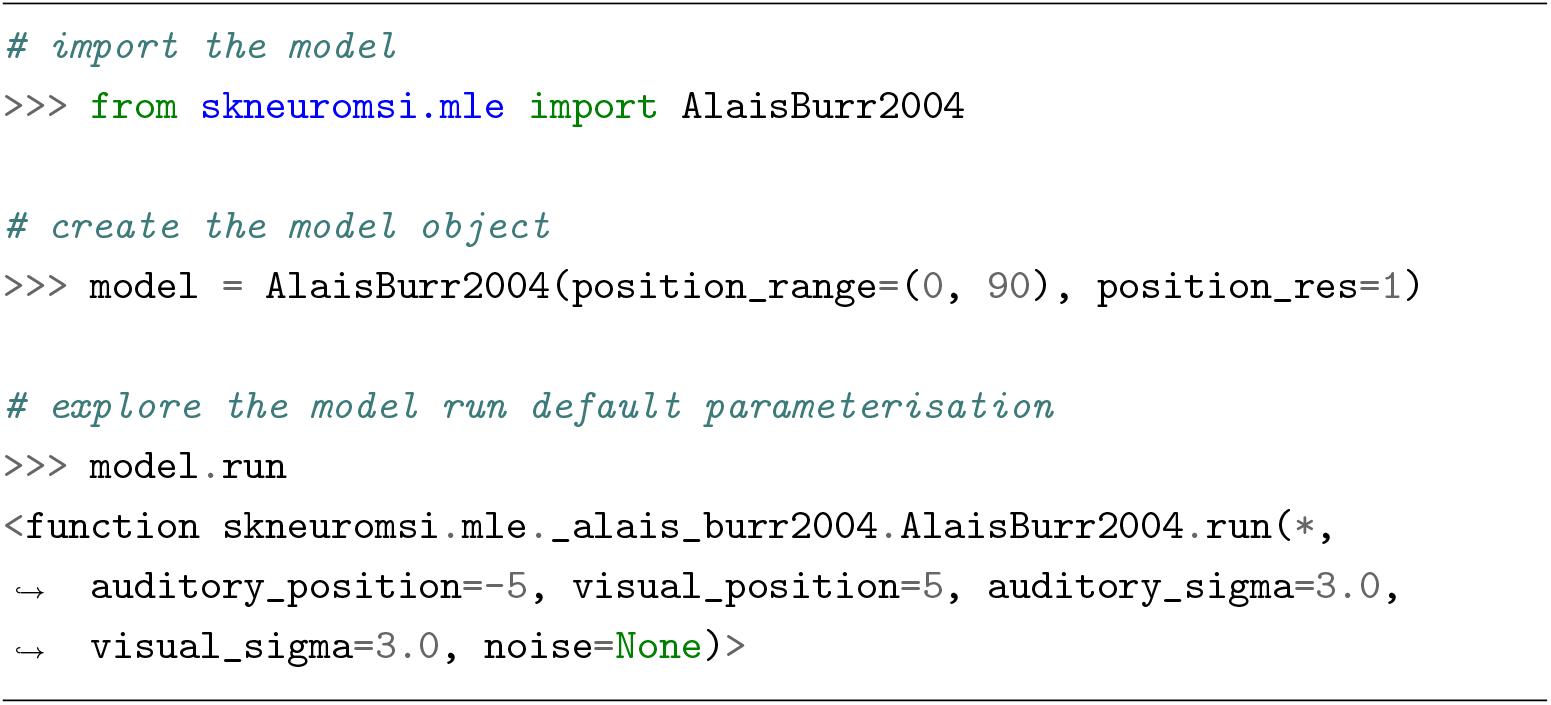
Example of Near-optimal Bimodal Integrator instantiation and default execution parameterisation within the Scikit-NeuroMSI package.

### C.2 Bayesian Causal Inference

The Bayesian Causal Inference model (Körding et al., 2007) uses the following formulation:

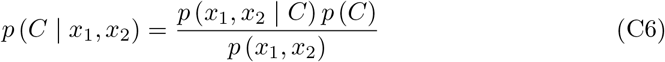

where *x*_1_ and *x*_2_ are two unimodal signals and C is a binary variable that represents the number of causes in the environment.

The posterior probability of the signals having a single cause in the environment is defined as follows:

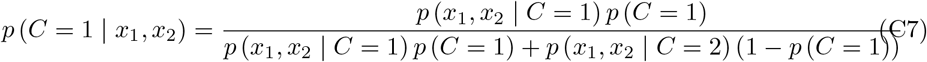

and the likelihood is computed as:

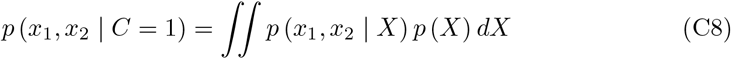

Here *p* (*C* = 1) is the prior probability of a common cause (by default 0.5). X denotes the attributes of the stimuli (e.g. distance), which are then represented in the nervous system as *x*_1_ and *x*_2_.

These equations show that the inference of a common cause of two unisensory signals is computed by combining the likelihood and prior of signals having a common cause. A higher likelihood occurs if the two unisensory signals are similar, which in turn increases the probability of inferring that the signals have a common cause.

The Bayesian Causal Inference model can be called using Code 8.

### C.3 Audio-visual integration and causal inference network

The audio-visual integration and causal inference network (Cuppini et al., 2017) consists of three layers: two encode auditory and visual stimuli, separately, and connect to a multisensory layer where causal inference is computed. Each of these layers consists of 180 neurons arranged topologically to encode a 180° space. In this way, each neuron encodes 1° of space and neurons close to each other encode close spatial positions.

Each neuron will be indicated with a superscript *c* indicating a specific cortical area (a, v or m for the auditory, visual or multisensory area, respectively). Similarly, each neuron will have a subscript *j* referring to its spatial position within a given area. Neurons in each layer have a sigmoid activation function and first-order dynamics:

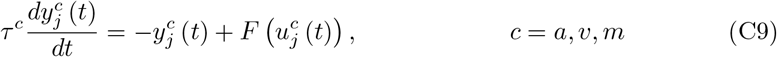

**Code 8.**
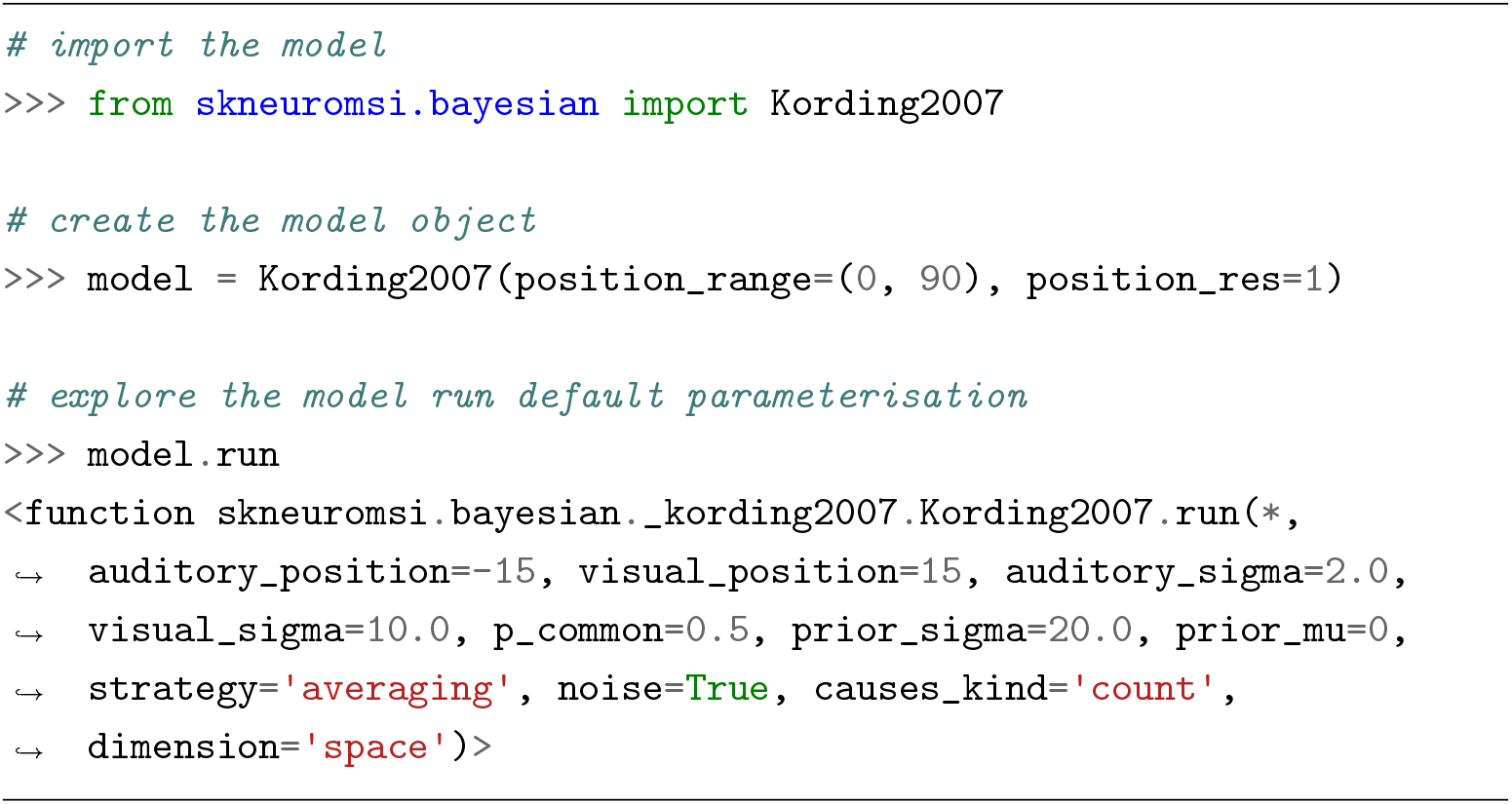
Example of Bayesian Causal Inference model instantiation and default execution parameterisation within in the Scikit-NeuroMSI package.

Here, *u*(*t*) and *y*(*t*) are used to represent the net input and output of a given neuron at time *t. τ*^*c*^ denotes the time constant of neurons belonging to a given area *c. F* (*u*) represents the sigmoidal relationship:

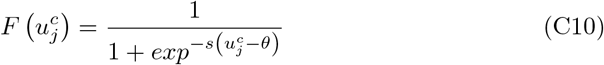

Here, *s* and *θ* denote the slope and the central position of the sigmoidal relationship, respectively. Neurons in all regions differ only in their time constants, chosen to mimic faster sensory processing for stimuli in the auditory region compared to visual stimuli.

These neurons are recurrently connected in a “Mexican hat” pattern within each layer. Such connectivity pattern consists of defining a central excitatory area surrounded by an inhibitory ring for each neuron, so that the entire layer generates excitation for spatially close stimuli and inhibition for distant stimuli:

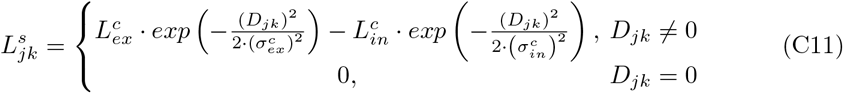

Here, 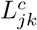 denotes the weight of the synapse from the pre-synaptic neuron at position *k* to post-synaptic neuron at position *j. D*_*jk*_ indicate the distance between the pre-synaptic neuron and the post-synaptic neurons within a given area:

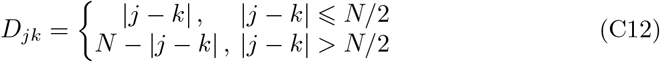

This defines a circular structure where each neuron receives the same number of lateral connections.

On the other hand, neurons in each unisensory layer (e.g. auditory) are reciprocally connected with neurons in the opposite layer (e.g. visual). These connections are excitatory and modify the spatial perception of unisensory stimuli. These synaptic weights are symmetrically defined 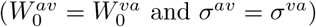 by the Gaussian function:

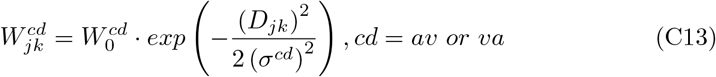

*W*_0_ denotes the highest level of synaptic efficacy and *D*_*jk*_ is the distance between neuron at position *j* in the post-synaptic unisensory region and the neuron at position *k* in the pre-synaptic unisensory region. *σ*^*cd*^ defines the width of the cross-modal synapses.

In addition, neurons in the unisensory layers have excitatory connections to the multisensory layer. These synapses are used to encode information about the mutual spatial coincidence of cross-modal stimuli and the probability that two stimuli were generated by a common source (i.e. they solve the problem of causal inference). The weights of these feedforward synapses are symmetrically defined as:

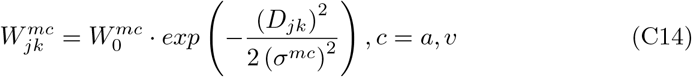

Here 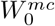 denotes the highest value of synaptic efficacy, *D*_*jk*_ the distance between the multisensory neuron at position *j* and the unisensory neuron at position *k*, and *σ*^*mc*^ the width of the feedforward synapses. The distance is defined as:

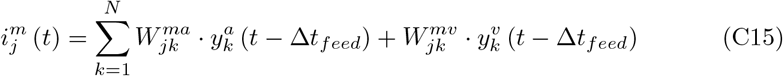

Here, Δ*t*_*feed*_ represents the latency of feedforward inputs between the unisensory and multisensory regions. 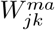 and 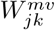 are the synapses connecting the pre-synaptic neuron at position *k* in a given unisensory area and the post-synaptic neuron at position *j* in the multisensory area.

Finally, the visual and auditory stimuli used as input to the network are defined with a Gaussian function to mimic the spatially localized external stimuli filtered by the receptive fields of the neurons. The stimulus from the external world is simulated as a 1-D Gaussian function to represent the uncertainty in the detection of external stimuli:

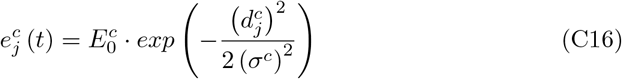

Here, 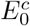 denotes the strength of the stimulus, 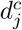 the distance between the neuron at position *j* and the stimulus at position *p*^*c*^, and *σ*^*c*^ the degree of uncertainty in sensory detection. The distance 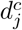 is defined as:

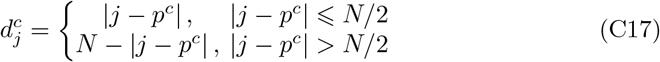

The central point of the Gaussian function corresponds to the point of application of the stimulus in the external world, while the standard deviation of the Gaussian function reflects the width of the receptive fields of the neurons and the reliability of the external input. This parameter is used to represent the different spatial acuities of the auditory and visual sensory modalities.

The net input of a neuron is the sum of an inside (i.e. within region) component 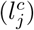 and an outside (i.e. extra-area) component 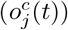:

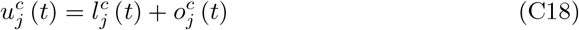

The within region component 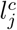 is defined as:

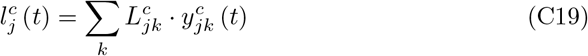

Here 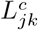 represents the strength of the lateral synapse from a presynaptic neuron at position *k* to a postsynaptic neuron at position *j* in the region *c*. 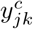 is the activity of the presynaptic neuron at position *k*.

Importantly, the extra-area input is defined differently for unisensory and multi-sensory areas. The extra-area input for the unisensory areas includes a stimulus from the external world 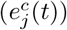, a cross-modal component coming from the other unisensory area 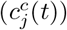 and a noise component 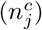. Furthermore, the cross-modal input is defined as:

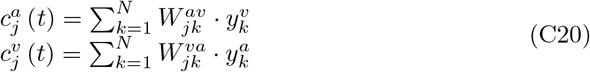

The noise component 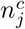 is extracted from a standard uniform distribution in the interval [*n*_*max*_ + *n*_*max*_]. Here, *n*_*max*_ is defined as the 40% of the strength of the external stimulus for each modality.

The Audio-visual Integration Network model can be called using Code 9:

### C.4 Multisensory Spatiotemporal Causal Inference network

The model consists of three layers: two encode auditory and visual stimuli separately and connect to a multisensory layer via feedforward and feedback synapses. At the unisensory areas, the model computes the spatiotemporal position of the external stimuli. In addition, at the multisensory area the model computes causal inference.

**Code 9.**
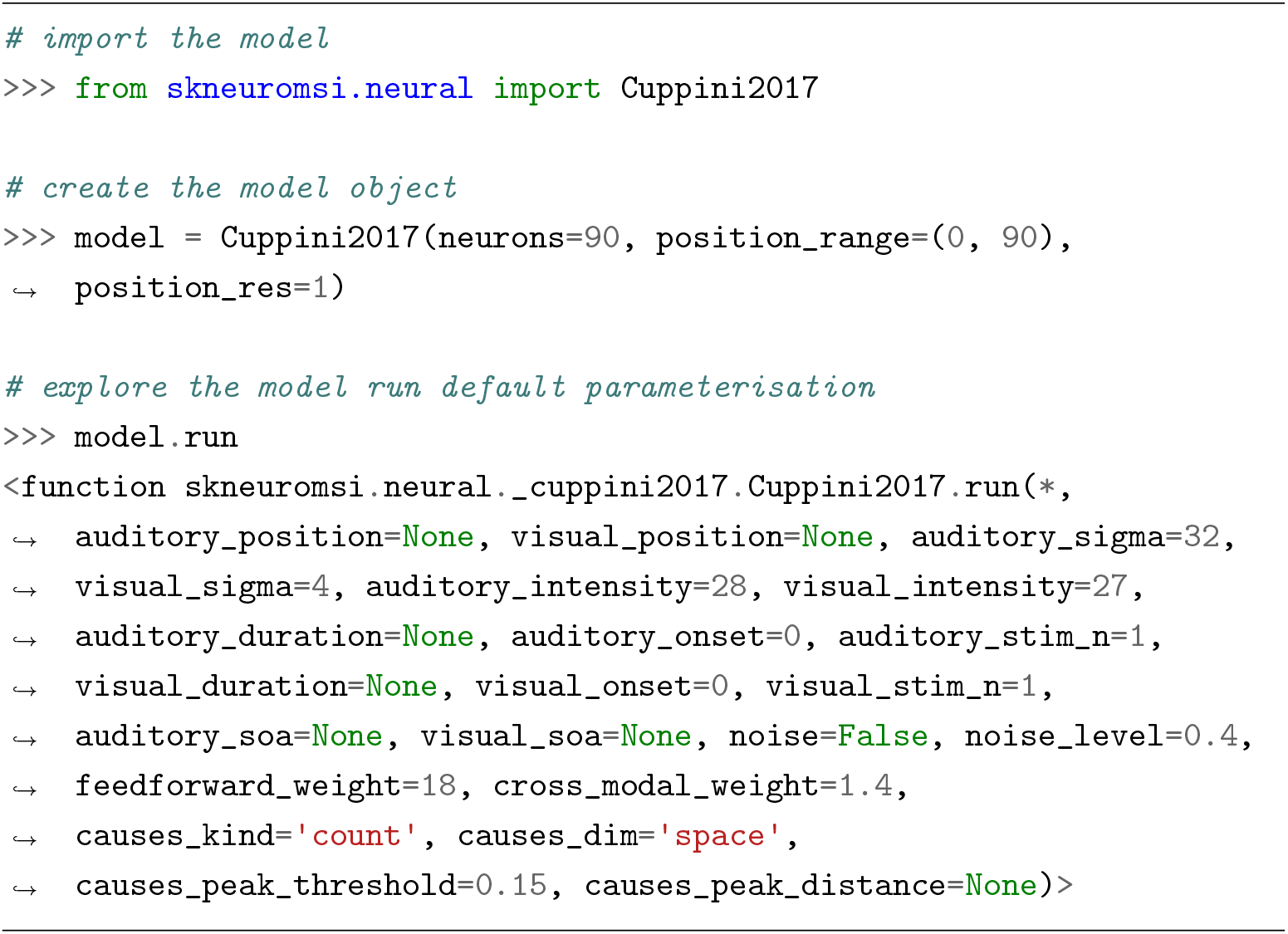
Example of the Audio-visual Integration Network model instantiation and default execution parameterisation within in the Scikit-NeuroMSI package.

This model maintains the neural connectivity (lateral, crossmodal, feedforward) and inputs described in the network presented in the previous section (Cuppini et al., 2017). Our model now includes feedback connectivity 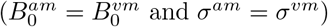, defined by the following equation:

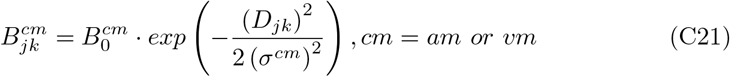

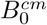 denotes the highest level of synaptic efficacy and *D*_*jk*_ is the distance between neuron at position *j* in the post-synaptic unisensory region and the neuron at position *k* in the pre-synaptic multisensory region. *σ*^*cd*^ defines the width of the feedback synapses.

Overall, the feedback input is defined as:

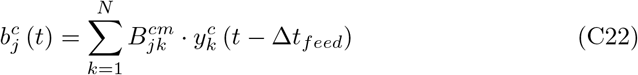

Here Δ*t*_*feed*_ represents the latency of feedback inputs between the multisensory and unisensory regions. The feedback synaptic weights are also symmetrically 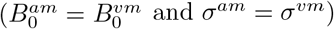 defined:

All these external sources are filtered by a second order differential equation to mimic the temporal dynamics of the stimuli in a cortex:

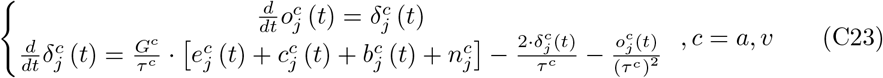

Here, *G*^*c*^ represents gain and *τ*^*c*^ the time constants of the dynamics.

These external sources are also filtered by a second order differential equation:

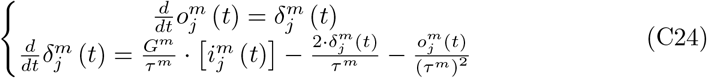

Here, *G*^*m*^ represents gain and *τ*^*m*^ the time constants of the dynamics in the multisensory neurons.

Furthermore, the cross-modal input is defined as:

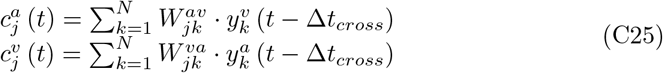

Here, Δ*t*_*cross*_ represents the latency of cross-modal inputs between two unisensory regions.

The Multisensory Spatiotemporal Causal Inference Network model can be called using Code 10.

**Code 10.**
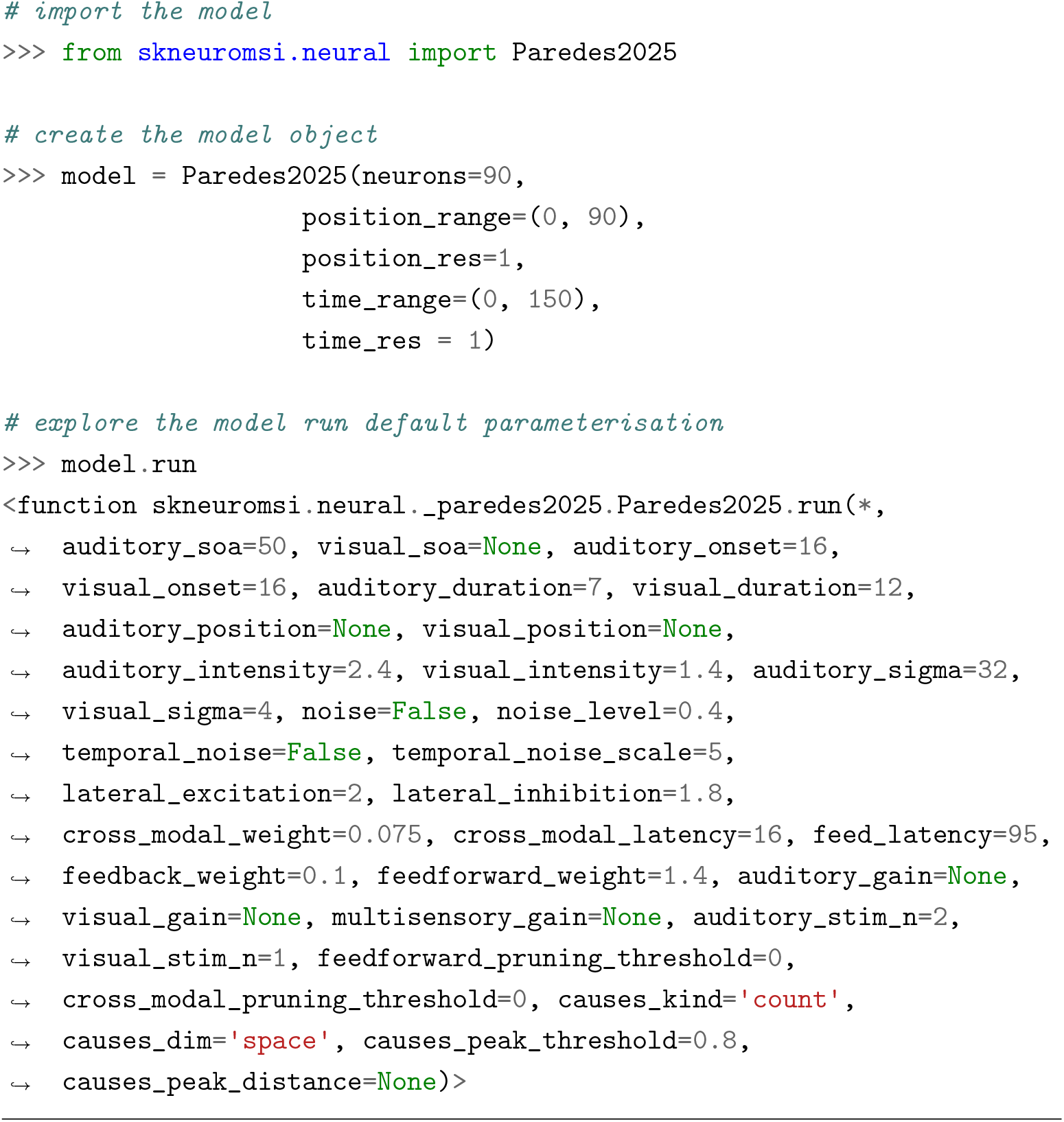
Example of the Multisensory Spatiotemporal Causal Inference Network model instantiation and default execution parameterisation within in the Scikit-NeuroMSI package.

## Appendix D Model fitting

### D.1 Model readouts

#### D.1.1 Implicit causal inference task

The audio-visual spatial localization task (Noel, Shivkumar, et al., 2022) involves participants determining whether an auditory stimulus is perceived to the right or left of a central position, indicated by a button press. Subsequently, psychometric functions are constructed by plotting the proportion of responses directed to the right as a function of the stimulus position. These data are modeled using a cumulative Gaussian function. The psychometric function provides two parameters that delineate the localization performance of the participants: bias and threshold. Bias is defined as the stimulus value at which responses are equally divided between rightward and leftward. A bias approximating 0° denotes highly accurate localization. The threshold is represented by the standard deviation of the fitted cumulative Gaussian function. A lower threshold indicates a higher precision in spatial localization.

In order to streamline the process and acknowledging the absence of noise in our simulations, we chose not to directly simulate the participants’ left-right responses. Instead, we concentrated on modeling the auditory position estimate for each model. The auditory position estimate for the network model was ascertained by pinpointing the neuron displaying the highest activity level during the simulation, with each neuron corresponding to a discrete spatial segment.

#### D.1.2 Explicit causal inference tasks

The explicit causal inference tasks require participants to ascertain whether auditory and visual stimuli originate from the same source. In the context of neural network models, this proportion is derived based on the maximal neural activation manifested within the multisensory neurons, contingent upon the condition that such activation exceeds the threshold of 0.15. In the spatial task, the evaluation of multisensory activity is conducted within the spatial domain at the concluding time point, while in the temporal task, the assessment occurs within the temporal domain at the locus of maximal activity. In cases where multiple peak values are identified, the average product of all potential combinations of these peak values is calculated to estimate the proportion attributable to the presence of multiple sources. Subsequently, the complementary proportion, representing the perception of multiple stimuli, is calculated to ascertain the proportion attributable to common source reports.

### D.2 Fitting procedure

The proportion of auditory bias or common cause reports reported by each model was fitted to the experimental data (Noel, Shivkumar, et al., 2022) with the cost function defined by Equation D26.

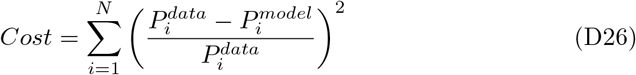

Here, 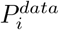 and 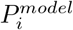 denote the proportion measured in the *i* th audio-visual stimuli disparity. N represents the number of disparities measured (i.e. 8 in the empirical study for the spatial tasks). This cost function was minimised by the implementation of the differential evolution algorithm (Storn & Price, 1997) available in the SciPy library for the Python programming language (Virtanen et al., 2020).

For the near-optimal bimodal integrator (Alais & Burr, 2004), parameters *σ*_*a*_ and *σ*_*v*_ were fitted under the constraint that parameter *σ*_*a*_ exceeds parameter *σ*_*v*_, thereby enhancing precision within the visual modality. The algorithm was provided with boundaries set at (0.1, 48) for both parameters.

For the Bayesian causal inference model (Körding et al., 2007), parameters *σ*_*a*_, *σ*_*v*_, *p*_*µ*_ and *p*_*σ*_ were also fitted under the constraint that parameter *σ*_*a*_ exceeds parameter *σ*_*v*_. For spatial tasks, the algorithm was provided with boundaries set at (0.1, 48) for all parameters, except *p*_*µ*_ which was set at (21, 69). For the temporal task, the algorithm was provided with boundaries set at (1, 500) for all parameters.

For spatial tasks, the audio-visual integrator and causal inference network (Cuppini et al., 2017), parameters *σ*^*a*^, *σ*^*v*^, *E*^*a*^ and *E*^*v*^, representing stimulus uncertainty and intensity, respectively, were fitted under the constraint that auditory uncertainty is larger than visual. The algorithm was provided with boundaries set at (0.1, 48) for uncertainties and (0.1, 30) for intensities. For the temporal task, the parameters 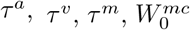, and 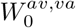 were fitted with boundaries set at (0.1, 55), (0.1, 55), (0.1, 75), (0.01, 150), and (0.01, 24), respectively. In the course of these simulations, the parameters *σ*^*a*^, *σ*^*v*^, *E*^*a*^, and *E*^*v*^ were consistently maintained at values of 32, 4, 50, and 49, respectively, with the duration of stimuli being fixed at 6 ms.

For the spatial tasks, the Multisensory Spatiotemporal Causal Inference network parameters *σ*^*a*^, *σ*^*v*^, *E*^*a*^ and *E*^*v*^ with the same boundaries and constraints as the previous model. For the temporal task, the parameters 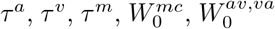 and 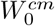 were fitted with boundaries set at (15, 50), (15, 50), (0.005, 50), (0, 7.5), (0, 0.15), (0, 0.005) respectively. During these simulations, the parameters *σ*^*a*^, *σ*^*v*^, *E*^*a*^, and *E*^*v*^ were consistently maintained at values of 32, 4, 2.55, and 2.5, respectively, with the duration of stimuli being fixed at 6 ms. Parameters Δ*t*_*cross*_ and Δ*t*_*feed*_ were assumed to be constant at 16 ms and 24 ms, respectively.

We acknowledge that employing a standardized fitting routine based on a genetic algorithm for noise-free model simulations may not constitute the optimal choice across all models. This approach may be particularly suboptimal for the Bayesian Causal Inference model, which was originally implemented utilizing a maximum-a-posteriori estimator with the assumption of random motor noise over 10,000 simulations (Körding et al., 2007).

## Footnotes

1 https://pypi.org/project/scikit-neuromsi/

2 https://scikit-neuromsi.readthedocs.io/

